# Single-Molecule Barcoding Technology for Single-Cell Genomics

**DOI:** 10.1101/2024.08.13.607508

**Authors:** Ivan Garcia-Bassets, Guoya Mo, Yu Xia, Tsai-Chin Wu, Immanuel Mekuria, Veronika Mikhaylova, Madison Rzepka, Tetsuya Kawamura, Peter L. Chang, Amber Paasch, Long Pham, Surya Shiv Venugopal, Sandra Sanchez, Janaina S. de Souza, Likun Yao, Sifeng Gu, Zsolt Bodai, Alexis C. Komor, Alysson R. Muotri, Joy Wang, Yong Wang, Ming Lei, Angels Almenar-Queralt, Zhoutao Chen

**Affiliations:** Universal Sequencing Technology Corp., Carlsbad (CA), USA; Department of Pediatrics, School of Medicine, University of California, San Diego, La Jolla (CA), USA; Sanford Consortium for Regenerative Medicine, La Jolla (CA), USA; Cellular & Molecular Medicine Department, School of Medicine, University of California, San Diego, La Jolla (CA) USA; Department of Chemistry and Biochemistry, University of California, San Diego, La Jolla (CA) USA; Universal Sequencing Technology Corp., Canton (MA), USA

**Author notes:** Correspondence to Z.C. (technology) or A.A.Q. (organoid biology). These authors contributed equally (Ivan Garcia-Bassets, Guoya Mo, Yu Xia).

## Abstract

Recent advances in barcoding technologies have significantly enhanced the scalability of single-cell genomic experiments. However, large-scale experiments are still rare due to high costs, complex logistics, and labor-intensive procedures. To facilitate the routine application of the largest scalability, it is critical to simplify the production and use of barcoding reagents. Here, we introduce AmpliDrop, a technology that initiates the barcoding process using a pool of inexpensive single-copy barcodes and integrates barcode multiplicity generation with tagging of cellular content into a single reaction driven by DNA polymerase during library preparation. The barcoding reactions are compartmentalized using an electronic pipette or a robotic or standalone liquid handling system. These innovations eliminate the need for barcoded beads and complex combinatorial indexing workflows and provide flexibility for a wide range of scales and tube formats, as well as compatibility with automation. We show that AmpliDrop is capable of capturing transcriptomes and chromatin accessibility, and it can also be adapted for user-customized applications, including antibody-based protein detection, bacterial or viral DNA detection, and CRISPR perturbations without dual guide RNA-expression vectors. We validated AmpliDrop by investigating the influence of short-term static culturing on cell composition in human forebrain organoids, revealing metabolic reprogramming in lineage progenitors.

## INTRODUCTION

Cataloging the vast diversity of cell identities and states in the human body and model organisms is a paramount scientific endeavor, foundational for understanding development, homeostasis, aging, and disease ^1^. A variety of single-cell genomic approaches, including single-cell RNA sequencing (scRNA-seq), have recently made this pursuit achievable, marking a new era in basic research with the promise to soon extend this unprecedented depth of high-content phenotypic characterization into translational and clinical applications ^2–5^. Over the past decade, the landscape of scRNA-seq technological innovation has rapidly evolved, and the many options currently available can be classified into three groups based on the underlying barcoding principle.

The first group applies bulk-like RNA-seq strategies to individually sorted cells in microwells (microliter-scale spaces),tagging the transcriptomic content from each cell with a distinct library index in every microwell ^6–12^. These methods process cells one by one, which largely limits scalability to no more than a few hundreds of cells in an experiment due to complex logistics and high costs. Still, these methods provide the highest sensitivity (gene capture) ^6–12^.

The second group achieves significantly higher scalability at a lower cost per cell by attaching millions of copies of a unique molecular identifier—a barcode—to a micron-sized bead and pairing beads with cells in thousands of nanoliter-scale spaces ^13–20^. These microscopic spaces can be created using water-in-oil droplets in an emulsion using a microfluidics instrument or an adapted vortexer in a templated emulsification; nanowells on multiwell plates; or a hydrogel, which exploits long polymers to limit diffusion in a tube without using physical barriers to separate the cells ^13–20^. Notably, all bead-based methods, except for 5’ versions, rely on reverse transcriptase as the barcoding enzyme, adding barcodes by priming the synthesis of first-strand cDNA with a bead-attached barcoded oligo-dT oligomer ^13–20^.

The third group of scRNA-seq methods further decreases costs per cell and can scale up to a million cells by exploiting the cells themselves as the compartmentalized spaces for the barcoding reactions ^21,22^. In these methods, cells are fixed and permeabilized and undergo three to four rounds of splitting and pooling on a 96-well plate, with ligase used as the barcoding enzyme, joining a well-specific index to first-strand cDNA in each split round ^21,22^. This process of combinatorial indexing randomizes the indexing at the cell level by providing a unique concatenation of three to four indexes to the transcriptomic content of each cell ^21,22^. For scaling to a million cells, combinatorial indexing on a 96-well plate is more userfriendly than preparing an equivalent number of libraries using a typical bead-based method. However, combinatorial indexing can be tedious when processing multiple sets of a million cells, and it can become impractical or less costeffective when processing only a few thousand cells.

Overall, there are three major deterrents to the popularization of large-scale scRNA-seq, aside from significant data analysis challenges ^23,24^: workflow complexity; inconvenient logistics; and, most importantly, high costs, since the limited gains in cost efficiency per cell can be offset by the opportunity to process more cells—Jevons Paradox.

Here, we report AmpliDrop, an innovative yet simple barcoding technology that skips the need for a combinatorial indexing scheme to achieve the largest scalability at a low cost per cell and the need for beads to introduce barcode multiplicity within a droplet. To achieve this, AmpliDrop introduces thermostable DNA polymerase as the third barcoding enzyme in single-cell experiments, enabling the generation of barcode multiplicity from a pool with millions of unique single-molecule barcodes during, rather than before, library preparation, unlike previous methods. Additionally, AmpliDrop offers flexibility through the use of a conventional electronic pipette or a typical robotic or standalone liquid handling system for the compartmentalization of the barcode multiplicity and barcoding reactions. These adaptations reduce the cost of barcodes to a negligible amount (a few cents per library), bring the cost of the barcoding reaction down to that of a PCR reaction, and allow flexibility for a broad range of reaction sizes—from a single library in a PCR tube to 96 independent libraries on a multiwell plate, with total throughputs spanning from a thousand to up to five million cells if the reactions are fully loaded, respectively.

## RESULTS

### Technology overview

The key elements of AmpliDrop applied to a 3’ scRNA-seq protocol are summarized in **Extended Data Fig. 1a**. The initial steps involve preparing the cells for subsequent in-cell enzymatic reactions. This preparation includes mild fixation to anchor RNA within cells and cell membrane permeabilization to allow reagent entry during the in-cell reactions.

The first in-cell reaction is reverse transcription (RT), where cells are incubated with reserve transcriptase and bead-free oligo-dT oligomers to synthesize first-strand cDNA (**Fig. 1a**, RT Step). The second in-cell reaction is RNA:cDNA tagmentation, where cells are incubated with Tn5 transposomes to fragment RNA:cDNA hybrids and introduce barcoding (BC’ing)-compatible sequences into the fragmented cDNA ends (**Fig. 1a**, Tagmentation). At this stage, cDNA-transposed cells are ready for encapsulation, lysis, and barcoding (**Extended Data Fig. 1b**, Part 1, Bulk Reactions).

**Fig. 1.**
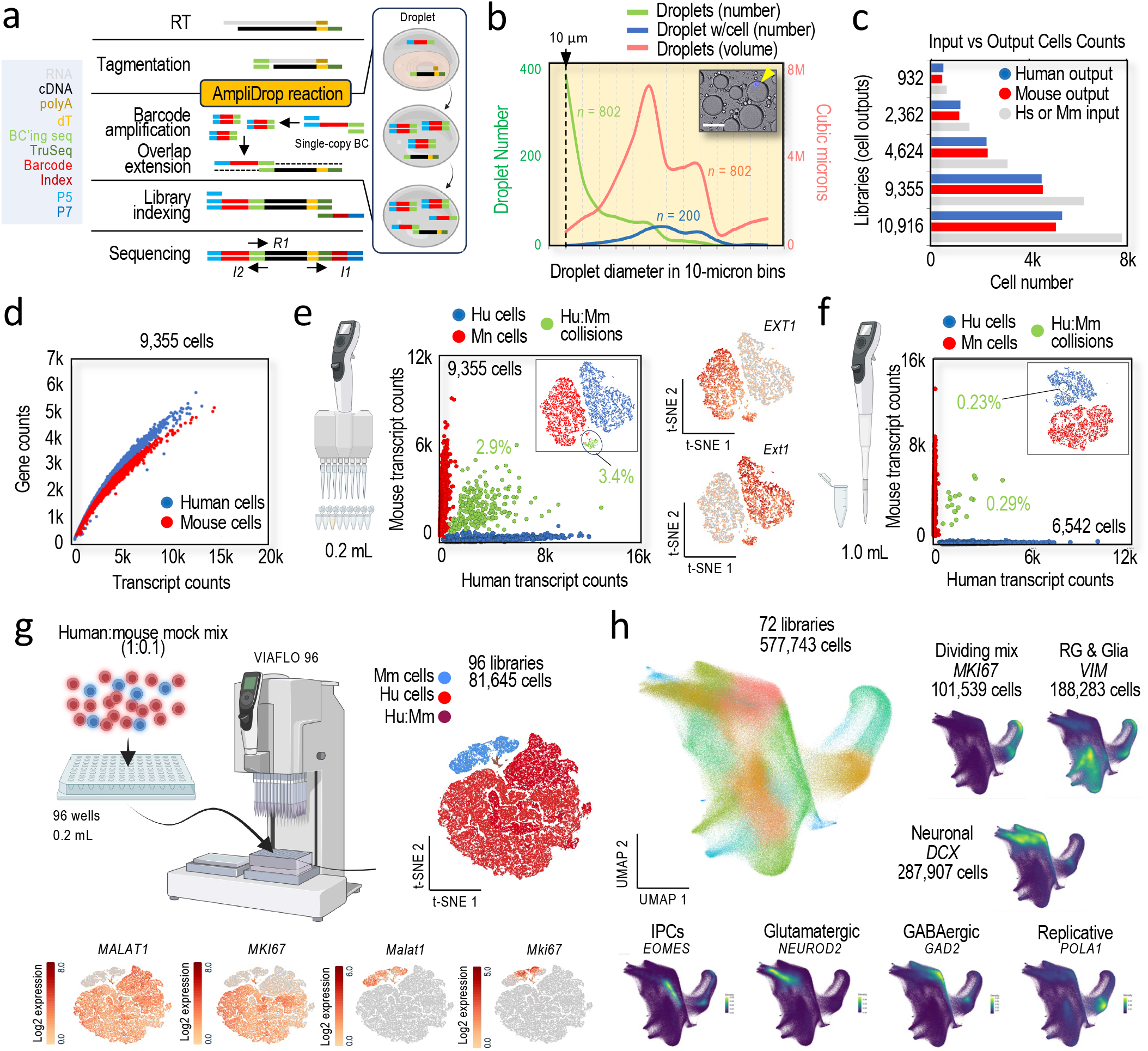
AmpliDrop barcoding applied to a 3’ scRNA-seq workflow. **a**, AmpliDrop library construction: Illumina Read 1 (R1) captures cDNA, Illumina Index 1 read (I1) captures the library index, and Illumina I2 read captures the cellular barcode. (Drawing) A unique single-molecule barcode is clonally amplified—i.e., within the droplet—and used to tag cDNA by overlap extension. **b**, ImageJ-based droplet number distribution in a 200 µL emulsion according to size, excluding n < 10 microns droplets (green line, left y-axis) and their estimated volume (red line, right y-axis) with data split in 10-micron bins (n = 6 microscopy images, n = 802 droplets). Included also, distribution of cell-containing droplets of any size (blue line, left y-axis; n = 142 microscopy images, n = 200 droplets). (Inset) Representative microscopy image depicting six droplets, including an estimated 54-micron droplet containing a DAPI-stained nucleus (arrowhead). Scale bar: 50 microns. Extrapolating the quantifications to the full emulsion, we estimated the generation of n = 804,548 droplets with a diameter larger than 20 microns, which should represent 99.5% of all the droplet-encapsulated cells and 98.07% of the total aqueous solution. **c**, Histogram shows human 293T and mouse NIH3T3 cell counts in five 200 µL emulsions (n = 5 libraries) pre and post encapsulation, pre-mixed at a 1:1 ratio. Pre-encapsulation counts (inputs) were estimated using the automated Countess Cell Counter (grey bars, same number for both species). Post-encapsulation counts (outputs) were estimated from the sequencing data according to read mapping behaviors (human in blue; mouse in red): 932, 2,362, 4,624, 9,355, and 10,916 cells. **d**, Scatter plot of transcript counts (x-axis) and gene counts (y-axis) by species in the 9,355-cell library (sequencing depth, n = 10,240 mean reads per cell). **e-f**, Scatter plots of transcript counts by species (human in x-axis and mouse in y-axis) in the 9,355-cell library with a 200 µL emulsion (in e) and the 6,542-cell library with a 1 mL emulsion (in f) (sequencing depth, n = 14,566 mean reads per cell). Cells color-coded by read mapping behavior (those separated from the axes were considered as human-mouse cell collisions in green). (tSNE plots) t-SNE plot insets in the scatter plots show color-coded cells by species based on gene expression. t-SNE plots on the right show expression levels (log2 scale) for the indicated genes (in e). Human-mouse cell collisions were inferred by their separate clustering behavior. **g**, (Top) Proof-of-concept 96 parallel 200 µL AmpliDrop reactions using 293T and NIH3T3 cells, pre-mixed at a 1:0.1 ratio, and encapsulated using a 96-multi-channel head (n = 81,645 cells, sequencing depth, n = 6,190 mean reads per cell). Cell identities and human-mouse cell collisions were color-coded based on human-mouse read behaviors. (Bottom) Expression levels for representative genes (log2 scale). **h**, AmpliDrop 3’ scRNA-seq analysis of 72 libraries and almost six hundred thousand cells. Cells were dissociated from a variety of human forebrain organoids (see Methods). Large UMAP plot shows data projected from 50,000 sketched cells (Seurat v5). Small plots show count densities for the indicated genes. Cell annotations typically associated with the indicated genes have also been included.

Next, a user-determined number of cDNA-transposed cells is encapsulated within droplets alongside PCR and barcoding reagents. These reagents include a pool of millions of unique single-copy barcodes, universal primers for their amplification, and a thermostable DNA polymerase. Encapsulation is achieved by mixing the cells and PCR and barcoding reagents with an emulsifying solution, using an electronic pipette or a robotic or standalone liquid handling system (for the largest scales). This process generates a highly thermostable emulsion (**Extended Fig. 1b**, Part 2, Single-Cell Reactions). The emulsion is then transferred to a thermocycler, where each encapsulated single-copy barcode is clonally amplified while the cells are lysed, releasing their transcriptomic content within the droplets (**Fig. 1a**, Barcode Amplification). In the same reaction, the already amplified barcodes are incorporated into cDNA by overlap extension, relying on the DNA polymerase again and the BC’ing-compatible sequences introduced by design at the 3’ end of the amplified barcodes and by random tagmentation at the 5’ end of cDNA (**Fig. 1a**, Overlap Extension).

The workflow ends with steps of droplet dissolution, barcoded cDNA pooling, and library indexing and amplification from the 5’ barcode side and the 3’ oligo-dT side, resulting in a 3’ scRNA-seq library (**Fig. 1a**, Library Amplification and Library Sequencing, and **Extended Data Fig. 1b**, Part 3, Library Preparation).

### Technology validation

To successfully work, AmpliDrop must minimize collisions— i.e., encapsulating two or more cells within the same droplet. We have optimized the mixing conditions to generate more than 800,000 cell-encapsulating-ready droplets in a 200 µL emulsion (**Fig. 1b**), which ensures at least 60 droplets per cell with loads lower than 15,000 cells (a high droplet-to-cell ratio).

To evaluate and validate these conditions, we performed a barnyard experiment using human 293T and mouse NIH3T3 cells pre-mixed at a 1:1 ratio. In a PCR-tube strip, using an 8-channel electronic pipette, we simultaneously prepared five libraries with inputs of 1,250, 3,120, 6,250, 12,500, and 15,600 cells. After sequencing, we recovered 75.28±1.19% of the cells (932, 2,362, 4,624, 9,355, and 10,916), largely preserving the 1:1 parity between species (**Fig. 1c**). Plots depicting ranked barcodes show a profile with the expected sudden drop in transcriptomic content, suggesting cell integrity with a high signal-to-noise ratio, or a robust cell-to-non-cell separation (**Extended Data Fig. 2a**). Most reads in the libraries, 83.32±5.07%, belong to cells, primarily mapping to the human or mouse genomes and transcriptomes at 84.78±0.15% and 74.22±0.31%, respectively. Moreover, using the 9,355-cell library as an example, we confirmed similar relationships between species in an analysis of gene capture by sequencing depth (**Fig. 1d** and **Extended Data Fig. 2b**). We also observed the anticipated segregation by species in a t-distributed stochastic neighbor embedding (t-SNE) plot (**Fig. 1e**, inset).

Regarding collisions, as expected, the fraction of estimated droplets with a mouse and human cell peaked with the largest outputs: 2.9% and 4.1%, equivalent to 5.8% and 8.2% collision rates, for the 9,355- and 10,916-cell libraries, respectively (**Fig. 1e**, 9,355-cell analysis). Importantly, these rates can be reduced without changing throughput by increasing the volume of the emulsion. In a 1 mL emulsion, for example, the inferred human-mouse cell collisions can be as low as 0.29% with an output of 6,542 mixed cells (**Fig. 1f**). Notably, the cost difference in PCR reagents between a 200 µL and 1 mL emulsion is less than $10, while the cell capacity is increased fivefold without changing the properties of the emulsion (i.e., same droplet-to-cell ratio and same droplet-to-barcode ratio). Throughput can also be increased by mixing multiple emulsions in parallel. For instance, mixing eight 200 µL emulsions in an 8-tube PCR strip using an 8-channel electronic pipette has a combined capacity for up to 80,000 cells: 8×10,000-cell emulsions. Likewise, simultaneously mixing ninety-six 200 µL emulsions using a 96-channel head on a 96-well plate has a combined capacity for close to a million cells: 96×10,000-cell emulsion (**Fig. 1g,h** and **Extended Data Fig. 2c**,**d**). Throughput can be further increased by simultaneously mixing ninety-six 1 mL emulsions in a 96-deep-well plate, with a combined capacity to process close to 5 million cells while still preserving the same encapsulating and barcoding properties as in a single 200 µL or 1 mL emulsion in 96 wells (**Extended Data Fig. 2e**). Data from many libraries can be then combined for an integrated analysis of a large number of cells (e.g., **Fig. 1h** and **Extended Data Fig. 3** show an analysis of 577,743 cells from 72 libraries).

### Inferring multi-barcoded cells

A challenge with encapsulation by pipette-mixing is achieving a one-to-one barcode-to-cell ratio. This requires using a low barcode-to-droplet ratio, which comes with the tradeoff of generating a high number of cell dropouts—droplets containing cells without a barcode. For example, according to a Poisson distribution ^25^, a barcode-to-droplet ratio of 0.01 should result in less than 1% multi-barcoded droplets but over 99% cell dropouts (**Extended Data Fig. 4a**).

To keep cell dropouts below 10%, we developed conditions that achieve an average barcode-to-droplet ratio of approximately three and created tools to computationally reconstruct multi-barcoded instances from the sequencing data (**Extended Data Fig. 4a**, line 3). In the aforementioned barnyard experiment with outputs of 932, 2,362, 4,624, 9,355, and 10,916 cells (**Fig. 1c**), these conditions resulted in an average barcode-to-droplet ratio of 3.22±0.32, with an estimate of 70% multi-barcoded cells and only 30% of cells with a single barcode (**Extended Data Fig. 4b**).

Reconstructing multi-barcoded cells requires an algorithm to match the transcriptomic partitions derived from the same cell based on read similarities. However, this process is challenging due to the sparsity of the scRNA-seq data and the expected abundance of cells with a similar transcriptome in any given cell mixture. To address these two issues, we leverage two features generating similarities among same-cell partitions.

The first feature is the diversity of 5’ ends generated by random Tn5 transpositions across the pool of cDNA molecules during in-cell tagmentation. We have termed the sequences at the transposition sites as ‘virtual unique molecular identifiers’ or vUMIs (**Fig. 1a**, tagmentation). The second feature is the possibility to generate some copies of the cDNA pool prior to barcoding. These copies can be generated by adding low amounts of poly-dT and BC’ing-compatible primers to the mix of PCR and barcoding reagents before encapsulation. Different copies of the same cDNA molecule (i.e., identical vUMI) can be then captured by the different barcodes within the same droplet. At a droplet scale, this should result in a distinctive vUMI pattern that could be used as a proxy of droplet origin. Notably, comparing vUMI and real UMI counts in pseudo-bulk analyses of the same data (the real UMIs were introduced in the oligo-dT primers) reveal a high correlation between both, thus suggesting that amplifying cDNA before barcoding does not generate obvious biases (**Extended Data Fig. 4c**).

To infer multi-barcoded cells, we have adapted an algorithm developed to identify multi-barcoded cells in scATAC-seq experiments ^26,27^, combining the reads from the barcodes that may capture transcriptomic signal from the same cell, while collapsing those reads with identical vUMI to eliminate potential pre-barcoding PCR duplicates from downstream analyses. We refer to the process of inferring a cell as ‘barcode merging’. The results, as shown in **Figure 1**, suggest the accuracy of the merging process. For example, the percentage of inferred human-mouse cell collisions remain relatively consistent pre- and post-merging: 4.47% and 3.40% (t-SNE-based) for the output of 9,355 cells, respectively (**Fig. 1e** and **Extended Data Fig. 4d**). We note that the slight increase in the pre-merging collision rate is likely due to the higher probability that collisions occur in larger droplets, which also contain a higher relative number of single-copy barcodes. Supporting this, the estimated collision rate in the 9,355-cell experiment is more than threefold higher in droplets with multiplicity of seven or more barcodes compared to those droplets with a barcode multiplicity of six or less.

Other results in **Figure 1** further support the accuracy of the barcode merging process. Specifically, the estimates of cell recoveries fall within the range observed with microfluid-based methods and largely preserve the 1:1 ratio between species (**Fig. 1c**). Additionally, collisions show high sensitivity to the volume of the emulsion, as expected from an accurate merging process (**Fig. 1c,f**). Despite these observations, we aimed to further validate the precision of the merging process by testing it with more challenging sample types.

First, we mixed four cell lines from the same species, human MCF-7, A2780, 293T, and HCT-116 cells, at ratios of 10,000:500:150:50, respectively, and compared the estimated and expected cell recoveries for four different outputs: 972, 2,539, 4,624, and 7,891 cells. Data visualization using uniform manifold approximation and projection (UMAP) revealed the expected four clusters in each case, which were annotated based on cell-line-specific markers identified in single-line libraries (**Fig. 2a**, plot, and **Extended Fig. 4e**). In support of accurate merging, the proportions of cells in each cluster largely matched the expected ratios across libraries, including for the cell line present at the lowest abundance (**Fig. 2a**, staggered plot).

**Fig. 2.**
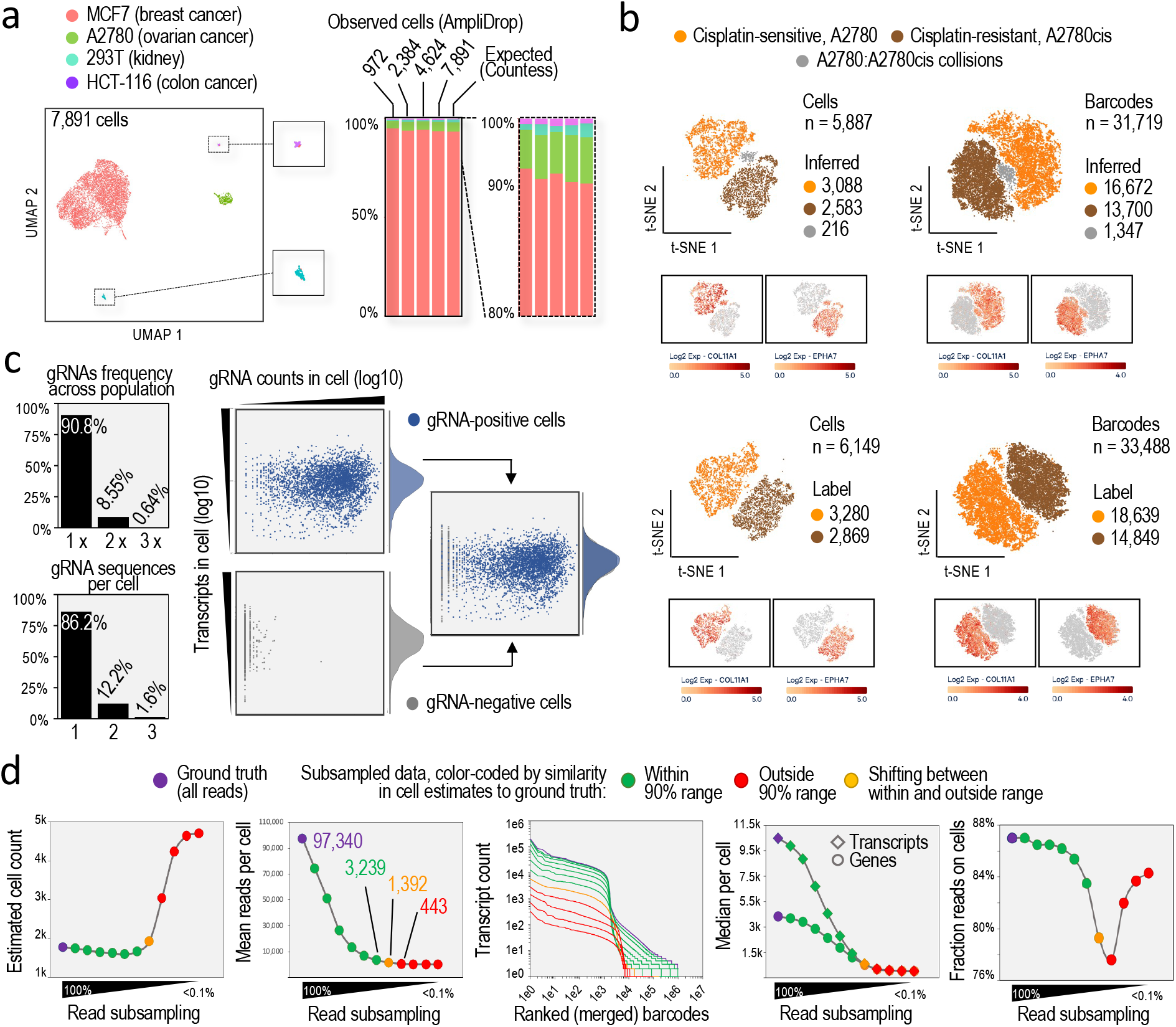
Reconstruction of multi-barcoded cell identities. **a**, (Left) UMAP plot of AmpliDrop 3’ scRNA-seq data based on human breast cancer MCF-7, ovarian cancer A2780, kidney 293T, and colorectal cancer HCT-116 cells, pre-mixed at a ratio of 10,000:500:150:50 (total, n = 7,891 cells; sequencing depth, n = 14,090 mean reads per cell). (Insets) Zoomed in images of the two smallest clusters. (Right) Stacked histograms show estimated cell proportions by predicted cell identity and output of cells. Cell identities were inferred based on markers validated in single-line experiments. A zoomed in version (scale: 80%-100%) is included for a better visualization. Stacked bars from Countess-based inputs (“ground truth”) have also been included. **b**, AmpliDrop 3’ scRNA-seq analysis (UMAP plots) based on drug-sensitive A2780 and drug-tolerant A2780cis cells, pre-mixed at a 1:1 ratio (top panels; n = 5,887 cells; sequencing depth, n = 12,542) and in silico mixed cells from independent A2780 and A2780cis libraries (bottom panels; n = 6,149 cells; sequencing depth, n = 11,608 mean reads per cell). In the pre-mixed sample (top panels), cell identities and A2780/A2780cis cell collisions (color-coded in grey) were inferred with markers from the in-silico mix (shown in the bottom panels). Left panels show plots with merged barcodes and right panels show plots with unmerged barcodes. **c**, (Left) Histograms of multimodal AmpliDrop 3’ and gRNA scRNA-seq data based on the gRNA-transduced K562 line (CRISPRi) show cell counts by the frequency of the gRNA sequence across the cell population (between one and three, top graph) or by the number of gRNA sequences per cell (between one and three, bottom graph). (Right) Scatter plots show cells by their amounts of gRNA counts (x-axis) and transcriptomic counts (y-axis), color-coded by the presence (blue) or absence (grey) of a high-confident gRNA sequence (left) or combined (right). Total, n= 1,340 gRNA-positive cells. Sequencing depth, n = 9,847 mean reads per cell (transcriptomic library) and 19,680,862 reads (gRNA library). **d**, AmpliDrop 3’ scRNA-seq analysis of Jurkat cells (n = 1,770) at high and subsampled sequencing depths (n = 97,340 mean reads per cell or ‘ground truth’, gradually subsampled to less than 0.1%). Panels (Left to right): cell counts, mean reads per cell, ranked barcodes by transcript counts, gene (circles) and transcript (diamonds) counts, and fraction of reads on cells. Results color-coded by similarity in cell-count estimates to ground truth, as follows: ground truth in purple, within 90% range in green, outside the 90% range in red, and transiting between within and outside range in yellow.

Second, we mixed two cell cultures of the same human cell line in two distinct cell states: one culture (A2780 cells) exhibiting high sensitivity to the anti-cancer drug cisplatin, and the other culture representing a drug-tolerant subpopulation (A2780cis cells), generated after repeated exposure and recovery from the drug ^28^. UMAP visualization shows clear segregation of the two cell states, based on expected markers ^28^, and nearly identical cell proportions pre- and post-merging: 43.88% and 43.19% for A2780, and 52.45% and 52.56% for A2780cis, respectively (**Fig. 2b**). The estimated collisions rates were also similar, 4.28% and 3.67%, with the pre-merging ratio slightly higher than post-merging ratio, as is also observed in the barnyard experiment (**Fig. 1e** and **Extended Data Fig. 4d**). Third, we assessed the accuracy of the merging process using a culture of lymphoblast K562 cells transduced with a library of 12,318-guide (g)RNAs at a multiplicity of infection (MOI) lower than 0.1 to limit the number of cells with more than one gRNA sequence. These sequences were inserted into the genome and used as an orthogonal molecular identifier (see Methods). During cell encapsulation, we included BC’ing-compatible primers against the gRNA flanking regions in the lentiviral construct. Thus, in every inferred multi-barcoded cell, all barcodes should capture the same amplified gRNA sequence if the merging process were accurate, since gRNA reads are excluded from the merging process.

As expected, data analysis reveal that most gRNA sequences can be found only once across the full set of inferred cells due to the high complexity of the gRNA library (**Fig. 2c**, left top panel). Also as expected, most inferred gRNA-positive cells contain only one high-confident gRNA sequence (**Fig. 2c**, left bottom panel). As an aside, we note that incorporating a gRNA readout into the AmpliDrop workflow did not impact the efficiency of the transcriptomic capture (**Fig. 2c**, scatter plots). About the accuracy of the merging process, we focused on the 3,159 barcodes assigned to the 1,838 inferred multi-barcoded cells with gRNA counts above the lowest 10^th^ percentile, avoiding ambiguities associated with low gRNA detection. Remarkably, all but one of the 3,159 barcodes (0.032%) were correctly assigned to a multi-barcoded cell where all barcodes capture the same gRNA sequence, representing 858 cells out of the total of 859 that were inferred. According to this estimate, only one out of 859 multi-barcoded cells would be incorrectly merged (0.116% error rate), which is lower than the estimated collision rate for a sample with a similar cell number. We note that the other barcode incorrectly merged in the same multi-barcoded cell was missed due to insufficient gRNA counts (below the 10^th^ percentile, further in support of its incorrect merging.

Finally, we evaluated the robustness of the merging process across a broad range of sequencing depths using a deeply sequenced AmpliDrop library prepared from Jurkat cells (1,770 cells; 97,340 mean reads per cell). We iteratively subsampled the reads from a 100% to as low as 0.0977% and subsequently merged the data on each subsampled dataset. The cell estimates, which serve as a proxy for correct merging, remained largely consistent until the average reads per cell dropped to 1,400, a point significantly lower than the typical sequencing depth reported in the scRNA-seq literature (**Fig. 2d**).

Together, these results support the robustness of our barcode merging strategy in inferring multi-barcoded cells, even with shallow sequencing depths and homogenous cell mixtures.

### Technology benchmark

To benchmark AmpliDrop, we compared AmpliDrop and 10X Genomics v3.1 technologies using a dissociated sample of 7-month-old neural organoids. To grow these organoids, we followed a semi-guided protocol known for introducing rich diversity of cell identities and transitioning cell states ^29,30^, recapitulating early developmental stages of the human brain cortex ^29,31^. We dissociated the organoids and split the cells into two aliquots, each processed with AmpliDrop and 10X 3’ scRNA-seq technologies. After sequencing the four libraries in the same lane of a flow cell, we downsampled the data to standardize the average number of reads per cell to 35,443 and the number of cells to 7,000. Following data integration, we annotated fifteen cell type or states based on well-known brain cell markers (**Fig. 3a** and **Extended Data Fig. 5a**).

**Fig. 3.**
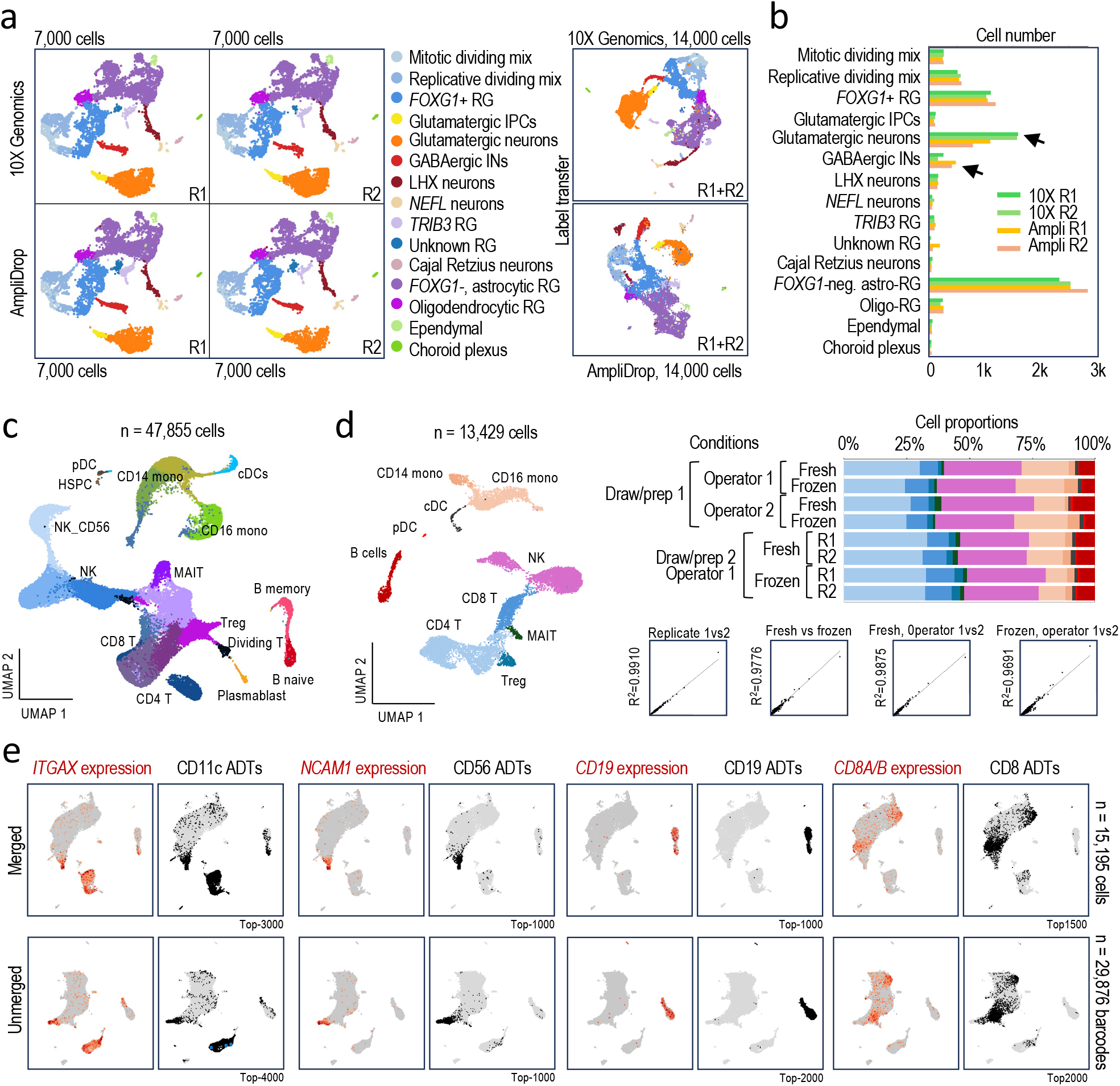
AmpliDrop 3’ scRNA-seq benchmarking. **a**, Comparative analysis between AmpliDrop and 10X Genomics v3.1 3’ scRNA-seq methods (two replicates each) using the same pool of dissociated human brain cells from 7-month-old cortical organoids. (Left) UMAP plots show integrated data from the four libraries standardized at n = 7,000 cells and n = 35,443 mean reads per cell (total, n = 28,000 cells) with data split by replicate and technology. Manual annotation based on well-established markers. (Right) UMAP plots of independently processed AmpliDrop (top) and 10X Genomics (bottom) data with annotations transferred from the full integration on the left. **b**, Cell count estimates color-coded by technical replicate and technology from the data shown in a. The arrows highlight consistent differences between technologies. **c**, UMAP plot of integrated PBMC AmpliDrop 3’ scRNA-seq data based on n = 47,855 cells (sequencing depth, n = 14,280 mean reads per cell). **d**, (Left panel) Comparative analysis of PBMC AmpliDrop 3’ scRNA-seq results based on n = 8 tests (total, n = 13,171 cells; sequencing depth, n = 8,571 mean reads per cell) to assess the technology robustness using a difficult sample type (outputs ranging between n = 869 and n = 3,014 cells): fresh versus cryopreserved, different operators, technical replicates, and different blood draws. (Top right) Stacked histograms show cell proportions by condition. (Bottom right) Scatter plots show gene-by-gene pseudo-bulk counts, as indicated conditions, with Pearson correlations. **e**, Multimodal cell-surface immunophenotypic and transcriptomic readouts (AmpliDrop CITE-seq) using TotalSeq-C TBNK-stained PBMCs (total, n = 15,195 cells; sequencing depth, n = 8,699 mean reads per cell). UMAP plots compare expression (log2 scale) and ADT (antigen-derived tag) counts (top cells based on the highest counts in black). as indicated. The bottom panels represent the same data (re-integrated) but pre-barcode merging.

Comparative analysis of cell identities and cell proportions shows high consistency between replicates and technologies across libraries. For instance (**Fig. 3b**), we detected virtually identical number of radial glia (RG) cells in the four libraries: 1,107-1,024 (10X) and 1,052-1,196 (AmpliDrop). Similar consistency was observed with replicative progenitors (514-572 and 542-585), mitotic progenitors (268-271 and 249-267), non-telencephalon neurons (168-160 and 168-153), and the small population of ependymal cells (67-58 and 53-56 cells). Some inconsistencies were observed between replicates in the smallest cell populations; however, these were not more prevalent in one technology over the other. This variability can partly be attributed to the challenge of capturing the diversity of a rich cellular mix in every subset of 7,000 cells. For example, while the number of *NEFL*-expressing neurons annotated with AmpliDrop was relatively consistent between replicates (66-57), the 10X replicates showed greater variation (62-100). Conversely, the number of choroid plexus cells was more consistent with 10X (47-49) than AmpliDrop (29-52). Two reproducible technology-associated differences were observed: a 1.68-fold enrichment in glutamatergic neurons in the 10X libraries relative to the AmpliDrop libraries, and a 2.09-fold relative enrichment in GABAergic interneurons in the AmpliDrop libraries relative to the 10X libraries (**Fig. 3b**, arrows).

Taken together, this side-by-side comparative analysis reveals that AmpliDrop and 10X methodologies capture similar cell diversity, both in terms of cell identities and cell proportions, with exception of two cell subtypes. Previous studies have also reported differences in cell proportions between single-cell technologies, suggesting that single-cell methods do not universally capture the same relative number of cells ^32^. Additionally, differential expression analysis shows that the 10X libraries are enriched in ribosomal and metabolic genes compared to the AmpliDrop libraries (**Extended Data Fig. 5b**,**c**, adjusted *p*-value < 10e-32 and Log2FoldChange > |2|). This finding aligns with previous reports indicating that 10X libraries disproportionally capture these gene classes compared to intrinsically nuclear-enriched methods, such as those based on combinatorial indexing, relying on fixed and permeabilized cells ^32,33^, like AmpliDrop.

To characterize cell diversity in a sample with well-established cell types, we prepared AmpliDrop libraries from 47,855 peripheral blood mononuclear cells (PBMCs), known for their low RNA content. We identified the expected repertoire of immune cells and cell states, including classical and non-classical monocytes, T and natural killer (NK) cells, B memory and naïve cells, mucosal-associated invariant T (MAIT), regulatory T cells, as well as rare cell populations, including hematopoietic stem and progenitor cells (HSPC), plasmacytoid dendritic cells (pDC), and plasmablasts (**Fig. 3c** and **Extended Data Fig. 6a**). We next applied unsupervised label transferring from a popular reference cell atlas to compare the cell annotations in our dataset with those from a publicly available experiment based on combinatorial indexing ^34^. This analysis shows that AmpliDrop and combinatorial indexing capture similar cell identities, although comparing cell proportions in this case would not be warranted since the PBMC samples derived from different donors (**Extended Data Fig. 6b**). We note that using combinatorial indexing to benchmark AmpliDrop is particularly impractical due to the need to accumulate enough number of samples to process a full 96-well plate.

To assess the robustness and reproducibility of the AmpliDrop data, we also used PBMCs due to their well-known fragility, comparing libraries prepared by different operators from the same or different blood draws from the same donor, as well as fresh versus cryopreserved cells, and performed technical replication. After data integration in a single UMAP plot (*n* = 15,151 cells), we observed the anticipated immune cell populations (**Fig. 3d**, UMAP, and **Extended Data Fig. 7a**). The libraries demonstrate consistency across replicates, operators, blood preparations, and fresh versus cryopreserved conditions (**Extended Data Fig. 7b**). In additional pseudo-bulk gene-by-gene analyses, technical replicates exhibit a correlation of r^2^ = 0.9910, fresh versus cryopreserved samples exhibit a correlation of r^2^ = 0.9776, and differences between operators exhibit a correlation of r^2^ = 0.9875 using fresh cells and 0.9691 using cryopreserved cells (**Fig. 3d**, scatter plots). Cell proportions also showed relatively similar estimates, although we expect some variation due to the small size of the libraries, which ranged between 869 and 3,014 cells (**Fig. 3d**, staggered bars).

As an alternative strategy to benchmark the accuracy of AmpliDrop in separating cell identities, we tested the technology for CITE-seq (cellular indexing of transcriptomes and epitopes by sequencing) ^35^. CITE-seq captures multimodal readouts, allowing comparison of cell identities and clustering behaviors using both transcriptomic and cell-surface immunophenotypic information. We stained a batch of PBMCs with a commercial cocktail of antibodies that react against nine immune-cell-surface antigens: CD19, CD3, CD16, CD4, CD11c, CD56, CD14, CD8, and CD45 (TotalSeq-C TBNK Cocktail). Each antibody is conjugated to a polyA-attached probe containing an identifier sequence compatible with AmpliDrop barcoding. After sequencing, probe-associated reads were excluded from the transcriptomic-based merging process and the subsequent steps of cell clustering and cell annotation (*n* = 19,958 cells; **Extended Data Fig. 8a**). Notably, probe capturing did not appear to affect the capture of the transcriptomic readout (**Extended Data Fig. 8b**). Comparing the transcriptomic and immunophenotypic readouts (both pre and post merging), we observed remarkably similar results, supporting data quality and, again, a robust process of barcode merging (**Fig. 3e**).

### Technology versatility

The value of any single-cell barcoding technology is enhanced with its versatility across different modalities and applications^36^. We have demonstrated that AmpliDrop can capture 3’ transcripts (**Fig. 1**), including in conjunction with genome-integrated gRNA sequences as part of a Perturb-seq lentiviral construct (**Fig. 2c**) and antibody-conjugated DNA probes (**Fig. 3e**). Next, we aimed to validate whether AmpliDrop could also capture other layers of cellular information.

The strong preference of the AmpliDrop 3’ scRNA-seq workflow to capture 3’ transcript ends results from adapter primers annealing to the 3’ end of the TruSeq-oligo-dT sequence during library amplification (**Fig. 1a**). Oligo-dT primers generally prime RT from terminal poly-A sequences in mRNA, except in those cases where oligo-dT primers anneal to intronic poly-A sequences in pre-mRNA. To more evenly incorporate non-3’-transcriptomic regions into the library, we added a second Tn5 activity to the tagmentation reaction, Tn5-ME-B, using primers that anneal to the second transposed sequence during library amplification (**Fig. 4a**, scheme, library 1). When applied to Jurkat cells, double Tn5A/B transposition increased the transcriptomic signal across gene bodies in normalized read density meta-profiles (**Fig. 4a** and **Expanded Data Fig. 9a**, reaction 1). The signal, however, remains partially biased towards the 3’ transcript end, suggesting that the RT reaction does not extend to the 5’ end in most transcripts. To confirm this hypothesis, a TruSeq-TSO primer (TruSeq-rGrGrG) was used to capture the 5’ transcript end. While the 5’ end of some genes, such as *ACTB*, was covered by this strategy, on a genome-wide scale, the shape of the meta-profile remained largely unchanged, thus, confirming an incomplete RT reaction for most transcripts (**Fig. 4a** and **Expanded Data Fig. 9a**, reaction 2). To increase the completeness of the RT reaction, we added random hexamers (R6) in combination with or without the TruSeq-oligo-dT primer during the RT reaction, which facilitated the coverage across gene bodies (**Fig.4a** and **Expanded Data Fig. 9a**, reactions 3 and 4, respectively). Together, these results reveal that AmpliDrop can successfully add barcodes with double Tn5A/B transpositions, enhancing coverage across transcripts (full-length).

**Fig. 4.**
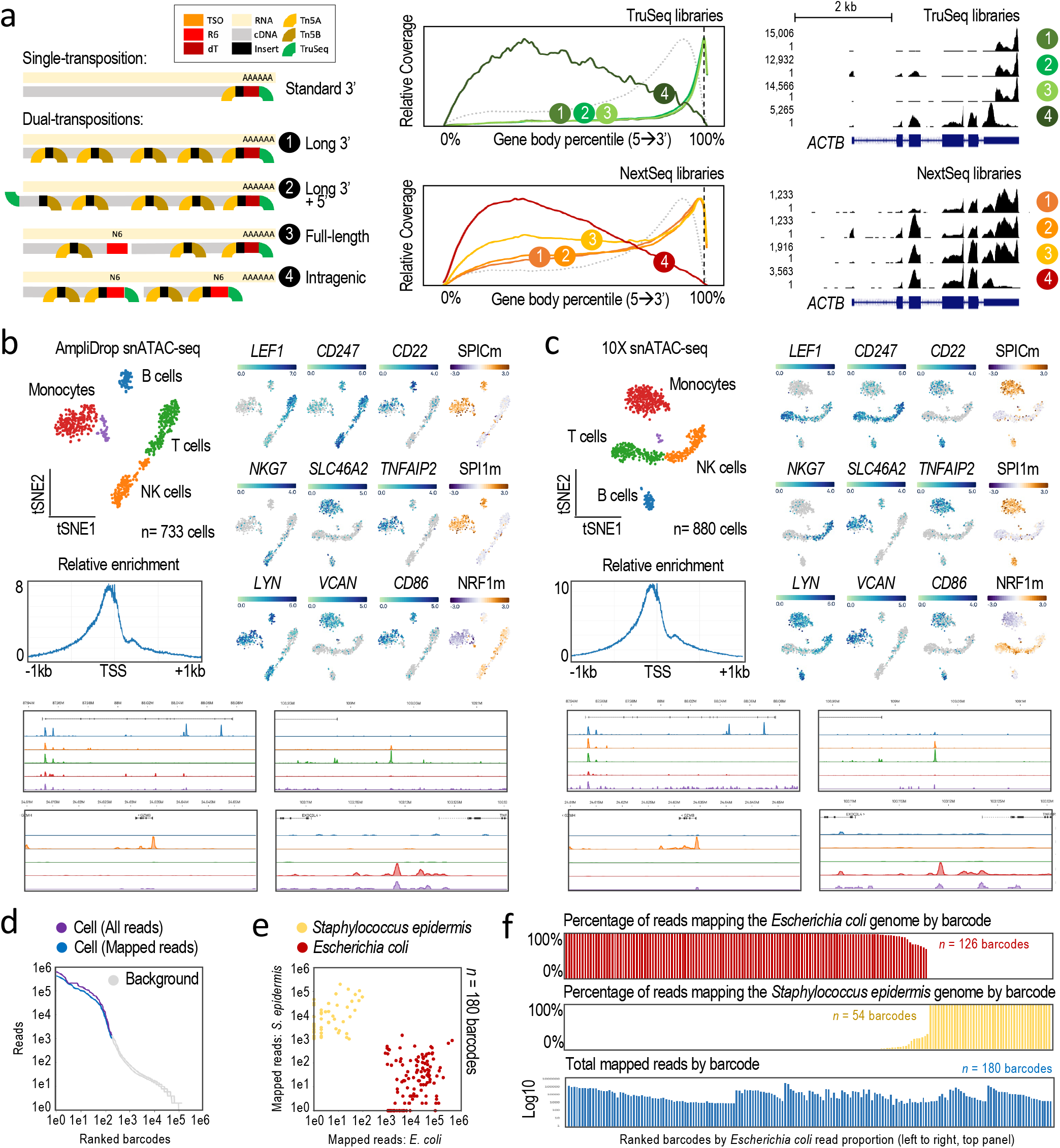
AmpliDrop barcoding applied to full-length scRNA-seq, snATAC-seq, and microbial scGenome-seq. **a**, (Left) Scheme represents AmpliDrop single (3’) and double (full-length) transpositions scRNA-seq strategies. (Center) Meta-profiles of AmpliDrop signal in Jurkat cells across normalized gene bodies, matching the numbers in the scheme. TruSeq (top) and NextSeq (bottom) libraries shown. For reference, the grey dotted line represents 10X 3’ scRNA-seq data (10k_hgmm_3p_nextgem_Chromium_X_Hu); the black dotted line indicates the submit of the AmpliDrop 3’ scRNA-seq signal in the TruSeq library. (Right) Read density across the ACTB locus, matching the numbers in the scheme. **b**,**c**, snATAC-seq comparative analysis with AmpliDrop PBMC data in b (total, n = 733 cells; sequencing depth, n = 9,533 mean reads per cell) and 10X Genomics PBMC data in c (total, n = 880 cells; sequencing depth, n = 8,718 median fragments per cell). The 10X Genomics data was previously generated and publicly available (atac_pbmc_1k_v1). Panels (clockwise): t-SNE plots with major PBMC annotations; t-SNE plots depict chromatin accessibility at the indicated promoter regions (blue gradients) and predicted enrichment of the indicated motifs in open chromatin regions (violet to brown gradients); tracks of pseudo-bulk read density by cell annotations across four representative loci, color-coded as in the t-SNE plot, and meta-profiles of snATAC-seq signal at and around TSS (1 kb on each side). **d**, Microbial scGenome-seq data: ranked barcode plots of AmpliDrop data from a mix of Escherichia coli and Staphylococcus epidermis cells with all or reference-mapped-only reads, as indicated (total estimated genome-capturing barcodes, n = 180). **e**, Scatter plot of reference-mapped read counts separated by barcode (each datapoint) and color-coded by species based on the highest read abundance (with n = 126 Escherichia coli and n = 54 Staphylococcus epidermis barcodes). Note: Reads with 0 counts were manually added to the log-scale plot. **f**, Histograms show the fraction of reads mapping to the Escherichia coli (top plot) and Staphylococcus epidermis (middle plot) genomes and the total number (bottom plot) by barcode (n = 180). Barcodes were sorted by the number of reads mapped to the Escherichia coli genome in all panels (decreasing left to right).

Using double Tn5A/B transpositions should also enable the capture of chromatin accessibility when following a single-nuclei (sn)ATAC-seq workflow ^37,38^. Notably, we observed no evidence of open chromatin signal crossover in double Tn5A/B-based scRNA-seq read density profiles (**Fig. 4a**, *ACTB*, and **Extended Data Fig. 9a**). To capture chromatin accessibility, therefore, we modified the library preparation, skipping the fixation and RT steps, isolating nuclei instead of permeabilizing cells, and amplifying and indexing the library with Nextera primers. To validate this protocol, we prepared four barnyard experiments based on human HCT-116 cells and mouse embryonic stem cells (mESCs) pre-mixed at a 1:1 ratio, estimating outputs of 505, 1,407, 4,312, and 12,298 nuclei. Multiple lines of evidence support the quality of the AmpliDrop scATAC-seq data, including (**Extended Data Fig. 10**): (i) a high similarity of read density profiles between pseudo-bulk and bulk data derived from our scATAC-seq and publicly available ATAC-seq data, respectively; (ii) similarity of collision rate estimates between AmpliDrop scATAC-seq and 3’ scRNA-seq data; (iii) the observation of a sudden drop in the cell-to-non-cell transition in ranked barcode plots; (iv) the observation of a meta-profile of read density showing the highest read accumulation upstream of the transcriptional stat site (TSS) ^39^; (v) the observation of a meta-profile of fragment size distribution that resembles a nucleosomal-like pattern of genomic DNA fragmentation, and (vi) the complete segregation of the human and mouse cells in t-SNE plots.

Next, to compare AmpliDrop and 10X snATAC-seq data, we generated an AmpliDrop snATAC-seq library from 733 PBMC nuclei extracted from a cryopreserved vial and compared its chromatin accessibility patterns and motif enrichment with a publicly available 10X snATAC-seq experiment based on 880 PBMC nuclei. Both were sequenced at a similar depth: approximately 42,500 and 40,700 average reads per cell, respectively. This comparative analysis shows similarities between the two methods, including comparable segregation of cell types in t-SNE plots, similar chromatin accessibility and motif enrichment across the promoters of immune cell markers, consistent read density accumulation upstream of TSS, and similar fragment size distribution with a nucleosomal-like pattern, among other key performance features (**Fig. 4b,c** and **Extended Data Fig. 11**).

We also applied double Tn5A/B transpositions to capture other sources of genomic information. In particular, the application of single-cell technology to microbial cell mixtures represents a promising advance in single-cell genomics ^40–43^. To interrogate the capture of microbial genomes with the AmpliDrop barcoding method—termed microbial scGenome-seq, we used a mix of gram-negative cells (*Escherichia coli*) and gram-positive cells (*Staphylococcus epidermis*) as starting material and prepared libraries following an snATAC-seq protocol with some adjustments primarily affecting the permeabilization step of bacterial cells (see Methods). A ranked barcode plot shows the expected signal drop, distinguishing cell-containing from empty droplets (**Fig. 4d**). Supporting single-cell behavior, most reads in the same barcode map either to the *Escherichia coli* or *Staphylococcus epidermis* genome, with an average coverage by barcode of 2.37% (126 barcodes) and 1.40% (54 barcodes), respectively (**Fig. 4e,f**). Aggregating all the reads from the barcodes by species (‘metacells’), we estimated a genome coverage of 90.68% and 30.05%, respectively (**Fig. 4e**), suggesting that AmpliDrop is capable of inferring pseudo-genomes in bacterial cell mixtures.

### Discovery potential

Finally, we aimed to validate AmpliDrop in addressing a biological question. Neural organoids are self-organizing multicellular structures used for modeling brain development and neurological disorders ^44^. However, it is often overlooked that these structures are fragile, susceptible to alterations caused by user handling, manipulation, and culturing ^45–48^. We used AmpliDrop to determine whether procedures included in some protocols, such as moving the organoids from the culturing plate to a secondary site for testing, keeping them outside the incubator during a testing period, or keeping them at least for short-term without shaking (static) during culturing can change cellular composition.

We split a pool of 7-month-old human induced pluripotent stem cell (hiPSC)-derived forebrain organoids into three groups. The first group remained in the incubator throughput the test. The second group was carefully transferred by aspiration to a secondary surface using a wide-orifice 1-mL tip pipette and returned to the culturing plate afterwards (**Fig. 5a**, transferred-only). The third group was processed as the organoids in the second group but was subjected to deformative compression to mimic an accidental damage during the transfer (**Fig. 5a**, transferred and damaged, and **Fig. 5b**, drawing). Once in the incubator, the two sets of manipulated organoids were maintained without orbital shaking to avoid disaggregation due to a potential higher fragility caused by their manipulation (especially with the third group). Two days later, the three sets were dissociated and processed for 3’ scRNA-seq.

**Fig. 5.**
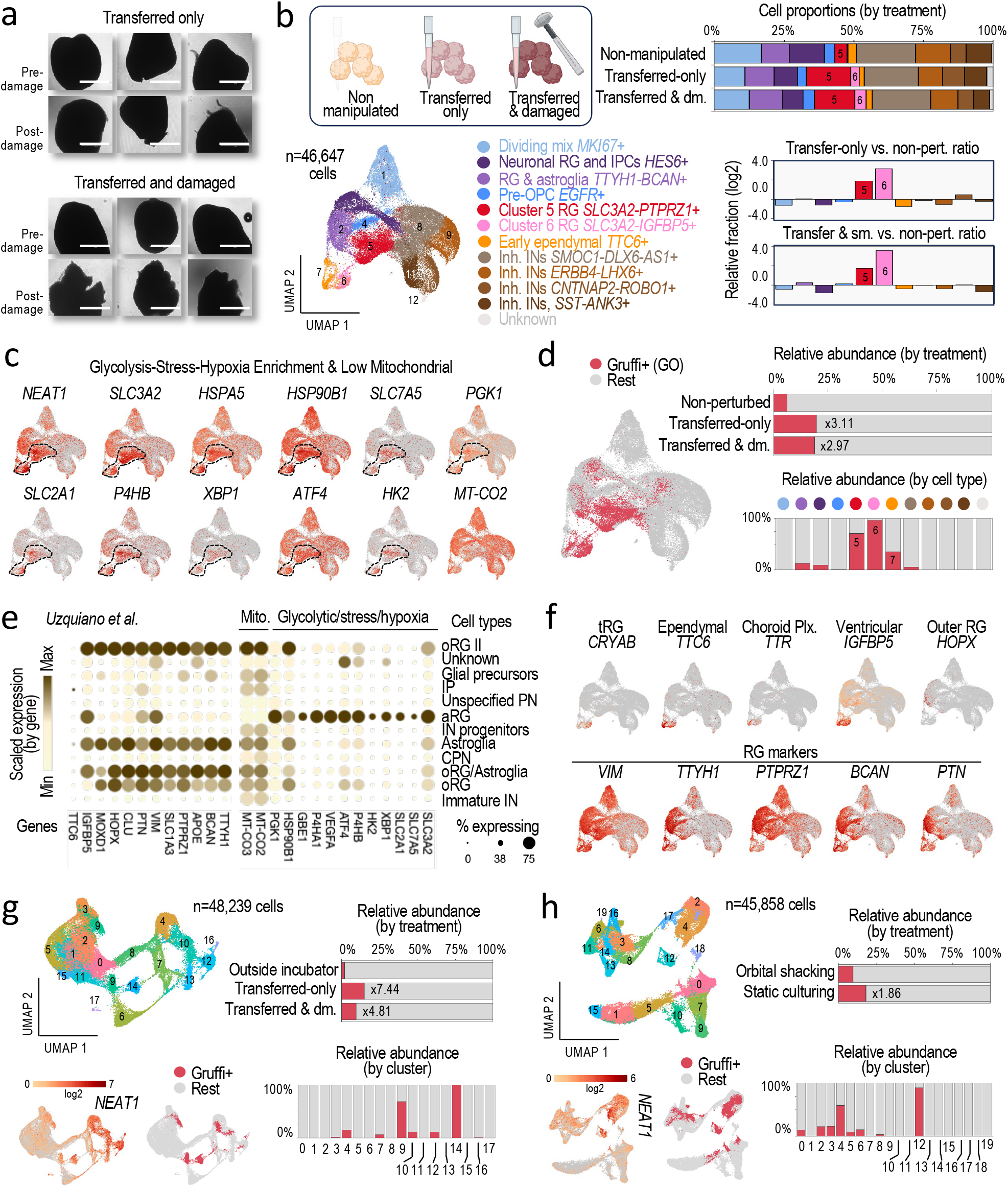
Influence of experimental factors on cell composition in human forebrain organoids: physical damage induced by transfer to a secondary site for potential experimentation (transferred-only and transferred-and-damaged to mimic an accidental deformation), time outside the incubator, and constant orbital shaking versus short-term static growth. **a**, Representative EVOS images show n = 7 seven-month-old forebrain organoids before and after manipulation. Scale bar: 1 mm. **b**, (Top left) Schematic representation of the three tested conditions. (Bottom left) AmpliDrop 3’ scRNA-seq analysis (UMAP plot) after integrating results from the three tested conditions, color-coded by inferred cell annotations (total, n = 46,647 cells; sequencing depth, n = 7,859 mean reads per cell). (Top right) Stacked histograms show cell proportions by test and cell type. (Bottom right) Cell proportions relative to the non-manipulated condition (log2 scale) show clusters 5 and 6 as the two most affected cell types/states. **c**, Expression levels (log2 scale) of glycolytic and stress/hypoxia-related genes on the UMAP plot shown in b. The dotted lines delineate the location of clusters 5 and 6, except for the mitochondrial gene, MT-CO2. **d**, (Left plot) Distribution of Gruffi-positive cells across the UMAP plot shown in b. (Top right) Histograms show counts of Gruffi-positive cells by condition. Numbers represent fold differences relative to the non-perturbed tests. (Top bottom) Histograms show counts of Gruffi-positive cells by cell type or state, color-coded as in b. **e**, Relative expression of genes distinctively expressed in clusters 5 and 6, including RG or glia markers, in the cell types annotated by Uzquiano et al., 2022 in their organoid data, visualized using the Single Cell Portal from the Broad Institute. **f**, Expression levels (log2 scale) of RG and glial markers on the UMAP plot shown in b, associated cell identities also indicated. **g**, (Top left) UMAP plot shows integrated AmpliDrop 3’ scRNA-seq data based on 6-month-old forebrain organoids in the indicated conditions, color-coded by clusters (total, n = 48,239 cells; sequencing depth, n = 9,760 mean reads per cell). (Top and bottom right) Distribution of Gruffi-positive cells by condition and cluster. (Bottom left) Expression NEAT1 levels (log2 scale) and Gruffi-positive cells across the UMAP plot. **h**, AmpliDrop 3’ scRNA-seq analysis of 2-month-old forebrain organoids grown in a static culture or orbital shaking for 3 days before cell dissociation. Panels organized as in g, color-coded by clusters (total, n = 45,858; sequencing depth, n = 17,209 mean reads per cell).

Notably, analysis of cell densities across the UMAP plot based on the integrated data (*n* = 46,647 cells) suggests significant changes in cell composition in the two groups of manipulated organoids (**Extended Data Fig. 12a**). Using a panel of brain cell markers, we concluded that these alterations mostly affected two subpopulations of radial glia (RG), clusters 5 and 6, characterized by the expression of the non-neuronal marker *VIM*, and advanced RG markers, such as *BCAN* and *PTN* (**Fig. 5b**, UMAP plot). Remarkably, the size of these two clusters increased by more than threefold and ninefold, respectively, compared to the non-manipulated set (**Fig. 5b**, graphs). We note that while cluster 5 shows a more classical RG-like identity, characterized by the expression of *PTPRZ1*, cluster 6 shows a more specialized identity, characterized by expression of a ventricular zone (VZ) glia marker, *IGFBP5*. Both clusters also show evidence of a distinct metabolic state, characterized by the increased expression of the mechano-sensing long non-coding (nc)RNA *NEAT1* ^49^, but not others mechano-sensing genes (such as *TRPM3, TRPC1, TRPV4*, or *YAP1*), and also the increased expression of glycolytic markers (such as *PGK1* and *HK2*) and hypoxia and cell stress genes (such as high *HSPA5, HSP90B1, XBP1*, and *P4HB* expression). Both clusters, furthermore, are characterized by low expression of mitochondrial genes (e.g., *MT-CO2*), without evident signs of mitochondrial or oxidative stress or apoptosis (low levels of *BCL2, BAX, SOD1, CASP8*, and *CASP9* expression) (**Fig. 5c** and **Extended Data Fig. 12b**).

In support of a distinct metabolic state for cells in clusters 5 and 6, analysis using a granular functional filtering approach, Gruffi ^50^, to locate cells enriched in genes associated with glycolysis and cell stress confirms that the two sets of manipulated organoids are enriched in glycolysis and cell stress gene ontologies by 3.11-fold and 2.97-fold, respectively (**Fig. 5d**, top). Gruffi-positive cells are mostly enriched in clusters 5 and 6 and, to a lesser extent, in cluster 7—a cluster that we annotated as ependymal cells due to selective expression of the ependymal marker *TTC6* (**Fig. 5d**, bottom).

Similar metabolically distinct cells have been recently reported in the inner core of organoids, annotated as apical-like RG (aRG-like cells) ^51^. In agreement, we confirmed that the same glycolytic and hypoxia-related genes enriched in clusters 5 and 6 were enriched in previously characterized aRG-like cells, which also show low mitochondrial gene expression (**Fig. 5e** and **Extended Data Fig. 12c**). Thus, we annotated cluster 5 as aRG-like cells and cluster 6 as tRG-like cells, the latter known to be a further differentiated RG subtype characterized by high *CRYAB* expression (as also observed in cluster 6; **Fig. 5f**, *CRYAB*) that ultimately evolves into ependymal and other glial cells ^52^. In the UMAP plot, cluster 6 cells are located adjacent to cluster 7 cells, which we annotated as ependymal cells (**Fig. 5f**, *CRYAB* and *TTC6*, and **Extended Data Fig. 13**).

Since the two sets of manipulated organoids were maintained outside the incubator for approximately 90 minutes (the time to take pictures pre- and post-manipulation and to transfer the organoids one by one from the culturing plates to the mental surface and back), we hypothesized that being outside the incubator, not the actual handling of the organoids, may have induced the observed changes in cell composition and metabolic state. To tests this, we conducted a second experiment with a set of 6-month-old organoids but, this time, the non-manipulated group was also kept outside the incubator as the other two groups (*n* = 48,239 cells after data integration). Still, we observed an increase in Gruffi-positive cells in the two manipulated conditions compared to the non-manipulated set: 7.44-fold and 4.81-fold, respectively (**Fig. 5g** and **Expanded Data Fig. 14**).

Next, we interrogated whether skipping orbital shaking after the manipulations might have been the underlying cause of the differences in aRG-like and tRG-like subpopulations. To test this possibility, we conducted a third experiment in which non-manipulated, younger organoids (2-month-old, which are richer in RG populations, were split into two groups, one group was maintained in the incubator without orbital shaking (static) for 3 days prior to scRNA-seq profiling, as in the previous two experiments, whereas the other group was maintained under orbital shaking for the same period. Remarkably, we observed an increase in Gruffi-positive cells in the absence of motion that mostly coincided with the *NEAT1*-positive subpopulation and primarily affected the central areas in the UMAP plot (clusters 4 and 12), where aRG-like and tRG-cells locate in the other two experiments, although in this case, cell proportions remained mostly unchanged (**Fig. 5h** and **Extended Data Fig. 15**).

Together, these analyses suggest that short-term culturing of the organoids without motion can change the metabolic state in some RG subpopulations, when damaging the organoids or maintaining them outside the incubator for 90 minutes, had not obvious effects. We cannot exclude the possibility, however, that the manipulations exacerbated the effects of culturing organoids without motion.

## DISCUSSION

We have developed AmpliDrop to address the need for greater scalability, simplified workflows, and streamlined logistics in single-cell genomic experiments. The core feature underlying these improvements is the generation of barcode multiplicity during, rather than before, library preparation, eliminating the need for costly barcoded beads ^13–20^ to isolate the copies of every unique barcode alongside cells and tedious combinatorial indexing schemes ^21,22^ to achieve large scalability. By generating barcode multiplicity during library preparation, furthermore, the source of barcodes (single-copy molecules) is largely negligible in a reaction while the cost of their amplification does not add any extra cost to the step of tagging (a PCR reaction). Importantly, the same pool of single-copy barcodes can be used for any number of libraries and throughputs, which simplifies the workflow. Additionally, using conventional multi-channel pipetting systems for encapsulating the barcoding reactions eases the logistics. Conveniently, furthermore, these systems (either electronic pipettes or automatic liquid handlers) are laboratory tools not exclusive for AmpliDrop use, in contrast to microfluidics devices or special vortexers used by other technologies.

Versatility without the need to substantially change the design of the barcoding step or reagents is another distinctive AmpliDrop feature. For example, most current single-cell barcoding methods require dual gRNA expression vectors to capture the gRNA sequence in a CRISPR perturbation experiment. With AmpliDrop, gRNA sequences can be captured directly by PCR from the gRNA-carrying DNA construct using user-customized primers added to the mix of reagents before encapsulation. The same principle can be applied to capture viral sequences (in infected cells) or bacterial or fungal sequences (inside single cells).

Regarding the versatility for using different tube formats, the most similar technology to AmpliDrop would be PIP-seq ^20^. However, there are some key differences between AmpliDrop and PIP-seq. In AmpliDrop, the barcode is the least expensive reagent, while in PIP-seq, bead-attached barcodes should carry a substantial manufacturing cost. AmpliDrop is compatible with robotic liquid handlers, facilitating automation, while this possibility is unclear for PIP-seq. It is also unclear the upper scalability limit for PIP-seq, although it is commercialized for a million-cell throughputs. While we have not generated 5-million-cell libraries with AmpliDrop due to excessive sequencing costs associated with this test, the 96-deep-well scalability is based on the same principle as those applied in a single PCR or 1.5 mL tube.

Another advantage provided by AmpliDrop is the generation of independent libraries on multiwell plates, unlike combinatorial indexing methods that generate a single multi-indexed library. This is relevant for the largest scalabilities. For example, for a CRISPR perturbation experiment at scale, AmpliDrop would generate ninety-six 50,000-cell libraries in a single 96-deep-weel plate (a total of 5 million cells). to enable the perturbation of all human genes with three distinct gRNAs for each gene (a total of 60,000 perturbations), aiming for 30 cells per gRNA sequence. Each library can be then sequenced separately until a number of sufficient hits is reached, facilitating the control of sequencing costs and data analysis. With combinatorial indexing, for example, twenty 96-well plates will be needed to process 5 million cells (or 4 set of 96-well plates for each group of a million cells), making more difficult the control of sequencing costs and data analysis, in addition to much more complex to process.

Proof-of-concept validation for AmpliDrop would not be complete without an example of its utility in addressing a biological question. We have focused on evaluating the susceptibility of neural organoids to alterations in cell composition induced by user manipulation. Neural organoids, which are valuable in vitro cell models for studying brain development and neurological disorders ^44^, are also extraordinarily fragile ^45–48^. This fragility makes them particularly vulnerable to handling and manipulation, potentially leading to confounding effects that can complicate data interpretation.

Supporting this vulnerability, previous studies have reported that mechanical forces generated during shaking—a common method for culturing these structures—can impact organoid structure and cell composition ^46^. Additionally, the speed of orbital shaking has been shown to influence both gross morphology and the microarchitecture of neural organoids ^48^. Conversely, it has been suggested that growing organoids under static conditions may lead to limited fluid dynamics of nutrients, negatively affecting the biology of these structures ^47^. Consequently, it remains unclear whether shaking or static growth represents a better system for modeling in vivo biology.

In this study, we examined the effects of various experimental procedures on cell composition in human forebrain organoids. After testing multiple hypotheses, our findings suggest that short-term static culturing, which is also used to generate assembloids ^63^, can increase the proportion of certain progenitor subpopulations, aRG and tRG, and/or enhance their glycolytic metabolism.

Currently, there is an ongoing debate in the neural organoid field regarding whether the expression of glycolytic markers in cell progenitors is a genuine characteristic of early brain development ^51,53,54^ or an artifact resulting from in vitro culture conditions, such as low oxygenation and poor nutrient access at the core of the organoid (where these cells locate). In the second case, these conditions may impair normal differentiation and should be, in a way, avoided ^50,55–62^.

Whether the effects observed in this study represent artifacts or true developmental properties—or a combination of both—remains unsolved. However, we find particularly interesting that previous studies suggest that shifting from aerobic glycolysis to mitochondrial oxidative phosphorylation (the opposite direction of the metabolic change that we would be observing, i.e., high expression of glycolytic genes and low expression of mitochondrial genes) is essential for neuronal differentiation, and that signs of cell stress would represent a homeostatic state in the early human brain ^54,64^. In fact, lncRNA *NEAT1*, which we find highly expressed in glycolytic aRG and tRG, is a major RNA moiety and scaffold of paraspeckles, and these structures serve as sensors of stress signals to allow cellular adaptation ^65^ and modulate differentiation and metabolism ^66–68^.

Thus, the question, remains whether replicating developmental biology be more accurate with orbital shaking or short-term static growth, i.e., with less or more glycolytic aRG and tRG. We speculate that during development, ventricular layer cells like aRG and tRG might act as sensors that use oxygen and nutrient levels as proxies for neocortex thickness (**Extended Data Fig. 13**). Given these cells’ potential to generate neuronal and glial populations in a time-controlled manner during development, a thin neocortex could signal differentiation into neuronal cell types early in development, while a thicker neocortex could signal differentiation into glial cell types later in development. We believe this metabolic property warrants further investigation.

In summary, this study based on organoids demonstrates that AmpliDrop can identify and characterize cell composition, as we have also recently reported in a study using assembloids ^69^.

## DATA AVAILABILITY

The data generated in this study has been deposited in NCBI GEO with accession number GSE272618. The following are the sources or accession numbers for data used in this study for comparative analysis: 65k PBMC Parse Biosciences scRNA-seq data from the site resources.parsebiosciences.com/downloads; 1k PBMC v1 10X Genomics snATAC-seq data from the site 10xgenomics.com/datasets; GSM5214800 for bulk HCT-116 snATAC-seq data, ENCODE data generated by the Mats Ljungman Lab (University of Michigan); GSM3399495 for bulk mESC snATAC-seq data ^70^, SCP1756 for the human cortical organoid data in the Single Cell Portal ^51^.

## ACKOWLEDGMENTS

Technology development and all proof-of-concept data was generated using private funding provided by Universal Sequencing Technology Corporation. The edited K562 line was generated through grant no. MCB-2048207, supported by the National Science Foundation to A.C.K. and the Research Corporation for Science Advancement through Cottrell fellowship no. 27975 to Z.B. The work on organoid biology was funded by the U.S. Department of Defense Peer Reviewed Alzheimer’s Research Program W81XWH-19-1-0315 and the Congressionally Directed Medical Research Program Traumatic Brain Injury and Psychological Health Research Program Idea Development Award W81XWH-21-TBIPHRP-IDA to A.A.-Q. The sequencing data for the organoid biology part was generated at the UC San Diego IGM Genomics Center. We thank Kristen Jepsen, Director of the UC San Diego IGM Genomics Center for help with sequencing utilizing an Illumina NovaSeq X Plus instrument that was purchased with funding from a National Institutes of Health SIG grant (#S10 OD026929). The formatting of this document is inspired by the submissions to bioRxiv from the Sergiu P. Pasca lab at Stanford University.

## CONTRIBUTIONS

I.G.B. and Z.C. (technology) and A.A.-Q. (organoid biology) conceived the study and supervised the research. G.M., T.-C.W, and I.M., (technology) and A.A.-Q. (organoid biology) performed most of the research. I.G.B., Y.X. performed most of the analyses. M.R., V.M., T.K., L.P., and A.P., (technology) and S.S.V. (organoid biology) contributed to the research. P.L.C., J.W., Y.W. contributed to the analyses. L.Y. (Fig.2b), S.G. Z.B., and A.C.K. (Fig.2c), and S.S., J.S.S., and A.M. (Fig.1h) generated reagents. I.G.B. wrote the manuscript (technology) also contributed by Z.C., G.M., Y.X., T.K., and A.P. A.A.-Q. wrote the manuscript (organoid biology). M.L., Z.C., and Y.W. gathered the funding to support the research (technology). A.A-Q. gathered the funding to support the research (organoid biology).

## METHODS

### Cell line culture

Human A2780 cells (ovarian, adenocarcinoma) and the cisplatin-resistant derivative A2780cis subclone were purchased from Sigma-Aldrich (Cat#93112519 and Cat#93112517) and cultured in RPMI-1640 Medium, GlutaMAX Supplement (ThermoFisher, Cat#61870036) supplemented with 10% fetal bovine serum (FBS, Omega Scientific, Cat#FB-11). Human MCF7 cells (breast, adenocarcinoma) were purchased from ATCC (Cat#HTB-22) and cultured in DMEM/F12 (ThermoFisher, Cat#11320033) supplemented with 10% fetal bovine serum (FBS, Omega Scientific, Cat#FB-11). The rest of cell lines were all also purchased from ATCC: human 293T cells (kidney, embryo, Cat#CRL-3216), human HCT-116 cells (colorectal, carcinoma; Cat#CCL-247), Jurkat cells (T lymphoblast, acute T cell leukemia; Cat#TIB-152), mouse NIH3T3 cells (fibroblast, embryo, Cat#CRL-1658). Mouse mESCs were a previously reported ^31^. All cells were grown in an incubator at 5% CO_2_ and 37°C, supplemented with penicillin-streptomycin (ThermoFisher, Cat#15140122), and sub-cultured as recommended by ATCC.

### Organoid culture

The WT83 clone6 of human induced-pluripotent stem cells (hiPSCs), derived from a typically developing Caucasian male ^29,71^, were grown on growth factor-reduced Matrigel (BD Biosciences, Cat#354234), coated 6-cm dishes in mTeSR Plus (STEMCELL Technologies, Cat#100-0276) without antibiotics. For propagation, cells were dissociated with Versene (ThermoFisher, Cat#15040066), 500 μM UltraPure™ EDTA (ThermoFisher, Cat#15575020), or ReLeSR™ (STEMCELL Technologies, Cat#100-0483). Cells were tested regularly for mycoplasma ^72^. To generate forebrain organoids, we followed a semi-guided protocol with a few modifications. Human iPSCs were grown in mTsER™ Plus (STEMCELL Technologies, Cat#100-0276) to a confluency of approximately 70% in a 6-cm plate and dissociated with a 1:1 mix of accutase (ThermoFisher, Cat#A1110501) /DPBS. Approximately, three million hiPSCs with a viability >95% were transferred into ultra-low attachment 6-well plates (Corning, Cat#3471) with mTsER™ Plus media supplemented with the SMAD inhibitors SB-431542 (Medchem, Cat# HY10431; 10 μm final) and Dorsomorphin (R&D Systems. Cat# 309310; 1 μm final), and the Rho-associated proteinase kinase (ROCK) inhibitor (RI) Y-27632 (Fisher Scientific, Cat# 125410; final 5 μm). Plates were placed on orbital shaker (95 rpm). Alternatively, hiPSCs were transferred to a well of AggreWell-800 plate (STEMCELL Technologies, Cat#34811) pre-coated with Anti-Adherent Solution (STEMCELL Technologies, Cat#07010) in mTeSR plus supplemented with RI. The AggreWell in mTsER™ Plus supplemented with RI and centrifuged to capture cells in microwells and transferred to incubator. The following day, embryoid bodies (EB) were transferred to a well of 6-well plate changing the media to fresh mTsER™ Plus supplemented with SB and Dorsomorphin as described above. From here, regular media and factors changes to complete generation of forebrain organoids were performed as reported ^30^, and maintained in M2 medium composed of Neurobasal (Life Technologies, Cat#21103049) supplemented with 1% GlutaMAX (Life Technologies, Cat#35050061), 1% MEM non-essential amino acids solution, NEAA (Gibco, Cat#11150-050), and 1x B27 (Life Technologies, Cat#17504044) performing half/medium changes every two days for the first month or twice a week after.

### Engineering and culturing of the gRNA lentiviral-transduced human K562 line

Wild-type K562 cells (ATCC, CCL-243) were grown in Roswell Park Memorial Institute (RPMI) 1640 Medium (Thermo Fisher, Cat#11875-119,) supplemented with 10% (v/v) fetal bovine serum and 1% penicillin/streptomycin (Gibco, Cat# 15140122). To construct the CRISPRi-expressing cell line, wild-type K562 cells were transduced with the dCas9-BFP-KRAB lentiviral vector (Addgene Cat#85969). Pure polyclonal populations of CRISPRi-expressing cells were then generated by sorting the transduced cells on a BD FACS Aria II instrument for the top 50% of BFP signal. The CRISPRi-expressing K562 line was then transduced with a gRNA lentiviral library containing 12,318 unique gRNA sequences adapted from hCRISPRi-v2 ^73^. Transductions were performed at a low multiplicity of infection (MOI < 0.1), using 2 × 10^8^ cells to ensure a representation of at least 500× coverage for each sgRNA after transduction. Transduced cells were enriched by treating with 2.5 μg/ml puromycin for 3 days. Cells were then cultured for 13 days before sequencing.

### Commercial and in-house isolated human PBMCs

For Fig. 3c,e, Fig. 4b, Extended Fig. 6, and Extended Fig. 7., cryopreserved human PBMCs were purchased from ALLCELLS (Cat#MNC, 10M). For Fig. 3d and Extended Fig. 7, peripheral venous blood was drawn from voluntary donor using EDTA-treated tubes and Lymphoprep™ (STEMCELL®, Cat. #29283-PIS 1 1 0) and stored at 4°C. Cells were resuspended in 1x phosphate-buffered saline buffer solution (1xPBS; Cat. #J61196.AP, Thermo-Fisher Scientific®) supplemented with 2% bovine serum albumin, BSA (Miltenyi, Cat#130-091-376), and layered over 1.5 mL Lymphoprep Density Gradient Medium (Stem Cell Technologies, Cat#07801) in a 15 mL conical tube, according to the manufacturer’s protocol. The solution was centrifuged at 800xg for 20 min at room temperature without brake. The mononuclear cell layer at the gradient interface was collected in a new tube and washed in PBS/2% FBS (10 mL total volume) and centrifuged at 120xg for 10 min at room temperature without brake. The wash step was repeated two more times, and the cells were resuspended in PBS for sequencing library preparation. The cells not used immediately were cryopreserved in 90% FBS/10% DMSO.

### Preparation of single-cell solutions (lines and PBMCs)

The following instructions detail how to dissociate adherent mammalian cells (optimized for 293T and NIH3T3 cells). Separately, grow cells in a 6-well plate according to standard procedures at 50-80% cell confluency (optimal), aiming to have 0.5-1.0 million cells from each line. Aspirate media and rinse twice with 1x PBS. Aspirate and discard the remaining 1x PBS. Add 150-300 mL of trypsin (ThermoFisher, Cat#25200056) and spread uniformly by tilting the plate. Incubate for 2-3 minutes at 37**°**C. When cells appear detached (move when shaking), inactivate trypsin by adding 1.3 mL of media supplemented with FBS. Aspirate and dispense the solution 5-10 times with a P1000 pipette against the surface to the plate to fully dissociate cells. Move cells to a 1.5 mL tube and centrifuge at 300xg for 5 minutes. Preferably, use a 4**°**C refrigerated centrifuges for all the spins. Resuspend in 1.5 mL 1x PBS, gently pipette up and down for five times, and spin down at 300xg for 5 minutes. Aspirate supernatant and gently resuspend in 1 mL of 1x PBS using a P1000 pipette at least 10 times for full cell dissociation. Take a 20 μL aliquot and mix with 20 μL of trypan blue (1:1) to determine cell viability using an automated cell counter, such as Countess (ThermoFisher), or manually with a Hemocytometer. Cells should appear isolated and cell viability should be at least 90% (unless a toxic treatment is applied to cells). If cells have been properly dissociated and show high viability, conditions are set to prepare cell solutions for AmpliDrop library preparation. For frozen PBMCs, cells were thawed in a 37°C water bath and, immediately, serially diluted with 1, 2, 4, and 8 mL of RPMI 1640 medium (Thermo-Fisher Scientific, Cat#11875093) supplemented with 10% fetal bovine serum (FBS; Sigma-Millipore) in the same 15 mL tube. While increasing the diluting volume, the tube was gently and manually rotated. Diluted cells were then centrifugated at 300xg for 5 min and gently resuspended serially in 1 mL and 9 mL of RPMI/10% FBS first prior to a new centrifugation. Cell pellets were gently resuspended in 1mL 1xPBS/0.4% BSA and, after a new centrifugation, resuspended again in 1mL 1xPBS/0.4% BSA before applying a 40 μL Flowmi Cell Strainer (Sigma Millipore, Cat#BAH136800040). Finally, cells were counted, and viability was examined using Countess III Automated Cell Counter (Thermo-Fisher Scientific).

### Preparation of single-cell suspensions (organoids)

For dissociation, 10-15 organoids were transferred into a 15 ml conical tube (BioPioneer, Cat#CNT-15). Media was aspirated leaving a small volume to avoid drying up. Organoids were rinsed twice with DPBS at room temp (3mL and 2mL), and cell dissociation was induced with 1.5-2 mL of a mix of StemPro Accutase (Life Technologies, Cat#A1110501), Papain (Worthington, Cat#LK003176), and DNAse (Worthington, Cat#LK003170) at a ratio of 2ml accutase:0.5 ml mix papain/DNAse prepared following manufacter’s instructions. The mix with organoids was kept static dissociating in the incubator at 37°C. Every 10 min until the organoids were dissociated (for a maximum of 50 minutes), organoids were gently but not slowly mixed 10 times with a 1 mL pipette tip/P1000 pipette. The dissociation was stopped by adding 4 mL of warm M2 media and pipetting with the P1000 pipette a few times. The solution was then filtered through a 40 µm filter (Falcon, Cat#352340), placed upside down on top of a 15 mL Precision™ conical tube (BioPioneer, Cat#CNT-15). The filter was first prewet with 1 mL of a 1 x DPBS supplemented with 0.1% BSA. Cells were harvested at 200xg for 7 min at room temperature. The media was aspirated leaving approximately 30 µL of supernatant. The cell pellets were resuspended in 0.7-1 mL of pre-chilled 1 x DPBS/0.1% BSA. From this point, all tips, tubes, and pipettes used were precoated with 1 x DPBS supplemented with 0.1% BSA to avoid cell loss. Gently, cells were mixed again by pipetting with a P1000 pipette and 120 µL of the solution was transferred to a 1.5 mL tube to count cells and assess cell viability using the ChemoMetec Cell Counter. Cell viability was above 90% in all cases (including for the severely damaged organoids). We then transferred around 500,000 cells to a new pre-chilled/pre-coated 1.5 mL tube and cells were centrifugated at 200xg for 7 min at 4**°**C, ready for immediate processing according to the AmpliDrop (or 10X) protocol.

### Organoid perturbation tests

We conducted three experiments. In the first experiment, two sets of 8 to 10 seven-month-old organoids were transferred from their original well in a six-well plate to a 24-well plate with a 10- or 25-ml pipette placing one organoid per well with fresh M2 media. A third set was maintained in the original plate in the incubator with gentle orbital shaking (88 rpm), as before separation. Pictures of the isolated organoids were taken with an EVOS microscope installed inside the safety hood. After taking the pictures, one set was returned to the incubator, and the other set was transferred using a wide-orifice (W-O) 1 mL tip onto a surface that is part of an in-home-built device that applies sudden compression to organoids (manuscript in preparation), inside the safety hood. After compression, the organoids were returned to the culturing plate by adding a few microliters of media around the organoids and aspirating with the 1 mL W-O tip again.

Organoids were never left to dry during this process. Post-compression, pictures were taken again with the EVOS microscope. The second set of organoids was processed as the first set but without mechanical compression. Afterwards, the cell culturing plate was returned to the incubator. In contrast to the non-manipulated set (third set), the other two sets were maintained without orbital shaking (static culturing) to avoid further damaging to the already mildly or severely damaged organoids. Two to three days later, the three sets were dissociated for scRNA-seq as described in the “Preparation of single-cell suspensions (organoids)” Methods section.

In the second experiment, three sets of six-month-old organoids were similarly processed as in the first experiment. The main difference is that the non-manipulated-organoid set was maintained outside the incubator as the other two sets, to eliminate differences caused by having the organoids outside the incubator (approximately 90 minutes, which results in media conditions changing as suggested by a transient media color change during this period). The organoids were finally dissociated as described in the “Preparation of single-cell suspensions (organoids)” Methods section.

In the third experiment, two-month-old organoids were transferred onto two wells in two different 6-well plates using a 50 mL pipette to minimize any physical damage. The plates were returned to the incubator for two days with one plate maintained under gentle orbital shaking (88 rpm) while the other plate was maintained in static culturing. Afterwards, the organoids were dissociated for scRNA-seq as described in the “Preparation of single-cell suspensions (organoids)” Methods section.

### AmpliDrop 3’ scRNA-seq

After automated cell counting and viability assessment using a Countess 3 Automated Cell Counter system (ThermoFisher), dissociated cells (typically, >90-95% viable) in 0.1% BSA-supplemented 1x DPBS buffer solution at this stage were processed with AmpliDrop 3’ scRNA-seq kits following the manufacturer’s recommendations (Cat#100050, 100051, 100052; Universal Sequencing Technology Corp.). Libraries were subject to quality control, sizing, and quantification using Agilent 4150 TapeStation system with High Sensitivity D1000 ScreenTape and Reagents. Libraries were sequenced on an Illumina NextSeq 500/550 instrument using a HighOutput kit (75 cycles) or on a NovaSeq 6000 or NovaSeq X instrument using a S4 kit (either PE50 or PE100) following the sequencing conditions: at least 51 cycles for Read1 + 8 cycles for Index1 + 20 cycles for Index2. In NovaSeq, we used 101×10×24×101 (PE100) or 51×10×24×51 (PE50) configurations used routinely in the Sequencing Core where we submit our samples with other customers. We noticed approximately only 5% lower reads per cell with the PE50 configuration compared to the PE100 configuration. In general, we processed only R1, I1, and I2, but when using oligo-dT primers with UMI, we also used R2 to capture the UMI sequence (10bp). Libraries were typically sequencing at a sequencing depth between 8,000 and 12,500 average reads per cell, unless otherwise indicated.

### Benchmarking AmpliDrop with 10X Genomics v.3.1 technology

Dissociated cells were split into two aliquots. One aliquot was immediately processed for 3’ scRNA-seq using the Chromium Controller system (10X Genomics) with a target recovery of 10,000 cells with the Next GEM Single Cell 3’ Reagent Kits v3.1 (Cat#1000268, 10X Genomics) and the other aliquot was also processed for 3’ scRNA-seq analysis by a second operator with AmpliDrop 3’ scRNA-seq (Cat#100050, 100051, 100052; Universal Sequencing Technology Corp.). In both cases, we followed the manufacturer’s recommendation for library preparation. Libraries were sequenced in the same lane at a 1:1 ratio with a S4 kit in NovaSeq. The AmpliDrop library was generated with UMIs incorporated into the oligo-dT primer, which added sequencing diversity in Read2 (10 bp), where the 10X library captures the insert. In Read1, the 10X library has sequence diversity provided by the UMI sequence (28 bp), where the AmpliDrop library has the insert.

For data comparison, the reads were down sampled to the same value as in the sample with the lowest reads among all samples, which was about 248.1 million. For the cells calling, we enforced to call 7,000 cells for each sample.

### AmpliDrop CITE-seq

PBMCs were stained with TotalSeq – A Human TBNK Cocktail (BioLengend, Cat#399901). One million PBMCs (typically > 95% viable) were resuspended in 45 µL cell staining buffer (BioLegend, Cat#420201), and 5 uL of Human TruStain FcX Fc Blocking reagent (BioLengend, Cat#422301) was added and incubated for 10 min at 4°C. TotalSeq antibody cocktail was reconstituted according to the manufacturer’s protocol and added to the blocked PBMC suspension and incubate for 30 min at 4°C. Cells were washed in 3 mL of cell staining buffer and centrifuged at 500xg for 5 min at 4°C three times. Cells were resuspended in 500 µL of cell staining buffer and filtered with 40 µm Flowmi cell strainer (Sigma, Cat#BAH136800040), and cell concentration and viability (typically > 90%) were recorded. The stained PBMC were immediately used as the input for AmpliDrop.

### AmpliDrop gRNA-seq (targeted AmpliDrop, Perturb-seq-compatible method)

To capture gRNA sequences, we followed the AmpliDrop 3’ scRNA-seq protocol with the following modifications. PCR primers were designed against the mouse U6 promoter and the gRNA backbone to amplify the protospacer sequence, targeting the amplicon size of approximately 140 bp. These primers were included in immediately before encapsulation at a final concentration of 50 nM. After the droplet breaking and the cleanup steps, half of the reaction was amplified with the Truseq index and P5 primer (3’ scRNA-seq), and the remaining half was amplified with Nextera index and P5 primer (gRNA sequencing). We sequenced the gRNA library with a NextSeq kit (150 cycles) as follows: R1, 70 cycles, R2, 70 cycles, I1, 8 cycles, and I2, 20 cycles.

### AmpliDrop Full-length scRNA-seq options

To explore full-length AmpliDrop options, we followed the 3’ scRNA-seq protocol with the following modifications. One million cells were used. We performed RT in the presence of switch oligo (TR2SW), or random hexamers (ProvTailedR6) with TruSeq sequences. Tagmentation was conducted with a mix of Tn5-A/B.

### AmpliDrop snATAC-seq

For snATAC-seq, we followed the AmpliDrop 3’ scRNA-seq protocol with the following modifications. Skipping the fixation step, cells were lysed in a modified permeabilization buffer (3.5 µl Permeabilizer in 100 µl Buffer P) and incubated on ice for 3-5 minutes, depending on the cell types (generally 5 min for cell lines, 3 min for PBMC) and 50,000 nuclei were used for tagmentation at 37°C for 60 min with a mix of Tn5-A/B. The cleanup step after the barcoding reaction was performed with 1.4x SPRI beads, and the Nextera index and P5 primer were used for the final ATAC library. The final cleanup was done with 1.2x SPRI beads.

### Microbial scGenome-seq

*Escherichia coli* DH10b And *Staphylococcus epidermis* FDA strain PCI 1200 (purchased from ATCC, Cat#12228) were grown in LB media at 37°C shacking at 200rpm. To capture bacterial sequences, we followed the AmpliDrop 3’ scRNA-seq protocol with the following modifications. *E*.*coli* BL21 (DE3) cells and *S*.*epidermis* cells were harvested at a late log phase, and 25 million cells (estimated by flow cytometry) from each culture were mixed at a 1:1 ratio. The mixed solution was washed once in 1 mL 1x PBS with 1 mg/mL probumin BSA (EMD Millipore, Cat#82-045-1), collected by centrifugation at 15,000xg for 5 min, resuspended in 50 µL of 1x PBS and fixed (AmpliDrop protocol). Following one round of wash as described above, the fixed cells were resuspended in 50 µL 1x PBS with 0.04% Tween-20 and incubated on ice for 3 min. Cells were then washed two times as above and incubated in 50 µL 20 mM Tris (pH8.0) containing 10 µg lysozyme (Thermo Fisher, Cat#90082) and 4 µg lysostaphin (Sigma-Aldrich, Cat#L7386-1MG) at 37°C for 30 min. The permeabilized cells were then washed two times as above, counted by flow cytometry, and 3 million cells were incubated in with tagmentation solution (AmpliDrop kit) containing double Tn5-A/B transposomes at 37°C for 60 min. The tagmented cells were washed twice and counted as above, and 2,000 cells were resuspended in 60 µL of the barcoding reagents (AmpliDrop kit). The barcoded products were recovered, amplified, and cleaned as in the 3’-scRNA-seq workflow. Libraries were sequenced on an Illumina MiSeq instrument using a Reagent Kit v3 (Cat#15043894, Cat#15043893, 150 cycles).

### Data analysis of gRNA sequences

When capturing PCR-amplified gRNA sequences incorporated into the AmpliDrop workflow, we generated a paired-end library to fully cover the gRNA design cassette with the gRNA sequences been flanking with the fixed lengths of sequences in R1 and R2. We detected gRNA sequences by matching the designed flanking sequences to the sequences showing in R1 and R2, with maximum of 3 hamming mismatches on each flanking side. We then counted any gRNA in each cellular barcode and filtered out the background noise with the gRNA less than 200 reads support or 2% of all detected gRNAs. The final gRNA counting table includes the detected gRNA sequence, the read supports, the percentage of this gRNA reads in all gRNA reads detected in the giving cell, and the unmerged and merged barcode sequences.

### Data analysis of CITE-seq DNA sequences

Antibody-conjugated probes were detected in the fastq files that contained cDNA reads. Briefly, we removed and saved the reads with their barcode information from fastq files if their sequences included any designed antibody-conjugated probe with exactly matching. We processed those fastq files without antibody probe reads with our pipeline so that the barcodes were appropriately merged and called cells. For each cellular barcode, we then counted the read numbers for each designed antibody probe from the file we saved as described above.

### Data analysis of AmpliDrop bacterial mock cell mixtures

We created the paired-end libraries for AmpliDrop bacterial mock cell mixtures. After demultiplexing, we mapped the paired-end reads with their barcode information to the reference genomes we used with bwa mem (0.7.17-r1188). We then parsed the bam files so that we could count the reads mapped the bacteria species within the giving barcode, and sorted based on the barcode ranks for all mapped reads. We plotted mapped reads with ranks, and we roughly considered the point that caused plot starting quickly dropping as the cell/non-cell boundary.

### Data processing for fastq subsampling

The deeply sequenced library from Jurkat cells was subsampled to 75%, 50%, 25%, 12.5%, 6.25%, 3.125%, 1.5625%, 0.7812%, 0.3906%, 0.1953%, and 0.0977%, respectively by randomly selecting reads from the original fastq file set.

### Generation of gene body percentile plots

We generated gene body percentage plots by sending the de-dupped bam files to the public tool of RSeQC (5.0.2) with the appropriate annotation (in .bed) file.

### AmpliDrop 3’ scRNA-seq data analysis

Sequencing data was processed using the AmpliDrop analysis software v1.0. Briefly, the software converts bcl sequencing files into fasq files using the Illumina tool bcl2fastq v2.20.0.422. Next, 8-nucleotide Illumina indexes (in I1) and 6-nucleotide AmpliDrop indexes (in I2) were error corrected for demultiplexing and separating reads for every library. RNA reads (R2) were trimmed with the adapter and poly-A sequences at the 3’ in the read using cutadapt v2.5. The bam contains the alignments with the paired end reads and contain the customized bam tag of the barcode for the aligned read. Reads were then aligned to the reference genome with STAR v2.7.10b in solo mode, ignoring mitochondria reads or those with a map score (MAPQ) lower than 30, and removing duplicates. Reads from merged barcodes were aggregated and those with identical vUMI were collapsed, eliminating potential PCR duplicates from downstream analyses. Fastq files with merged barcodes were then used as input files for count matrix generation using either Cell Ranger v5.0.1 (10X Genomics) for benchmarking purposes or Kallisto-bustools (kb). The first has been optimized for processing data from the Chromium platform, providing a solution that includes barcode processing, read alignment (using the STAR aligner), and quality-control metric, as well as the generation of popular output files, including the count matrix file in several formats. It is a very user-friendly tool. The second is an open-source option, which is more computationally efficient and fast, with pseudo-aligns reads to produce a barcode, UMI, set (BUS) file, then converted into a cell-by-gene count matrix ^77^. In all options, we included introns to quantify genes and UMIs. When merging was skipped, fastq files were processed skipping the merging step.

### Cell Ranger analysis for 10X 3’ scRNA-seq data

For the analysis of 10X 3’ scRNA-seq data, we used Cell Ranger v5.0.1 software (10X Genomics) following the developer’s instructions, selecting the option to include intronic reads --include-introns.

### AmpliDrop data integration using Seurat

In R Studio (v2023.03.0+386), we used the Seurat packages v4.1.1 ^34^ and v.5.0.1 ^24^ for data integration. We recommend reading about the impact of data package selection in scRNA-seq ^78^. Briefly, Seurat objects were created using the CreateSeuratObject() function from filtered_feature_bc_matrix files without applying filters other than min.cell = 3 and min.features = 200 to assess the quality of the AmpliDrop barcoding technology without computational aids. Moreover, in contrast to 10X technology, AmpliDrop is not characterized by an excess of mitochondrial and ribosomal signal capture and is less likely to introduce poor quality cells (so-called ‘dead’ cells) into the analysis, making this filtering much less necessary than with 10X technology. Next, the data was normalized with the NormalizeData() function using the LogNormalize method with a scale factor of 10,000. For PCA, we used gene expression variation detected with the FindVariableFeatures() function and the vst method, limiting to nfeatures = 2000. Then, the data was scaled up using the ScaleData() function for all.genes. We created a Seurat object for every sample. For integration, we used the FindIntegrationAnchors() and IntegrateData() functions with 20 dimensions. For multi-dimensional reduction and clustering, we used the ScaleData(), RunPCA(), RunUMAP(), FindNeighbors(), and FindClusters() functions with npcs = 30, reduction = “pca”, dims = 1:20, and a resolution = 0.5. To identify markers, we used the FindAllMarkers() function with logfc.threshold = 0.25. Data visualization was based on the ggplot2 v.3.5.1 library using the DimPlot() function. Projections were exported into csv files to import them into Excel v.16.84 (Microsoft).

### Comparative analysis between AmpliDrop snATAC-seq and 10X Genomics snATAC-seq data

10X Genomics data was obtained from the 10X Genomics website https://www.10xgenomics.com/datasets/1-k-peripheral-blood-mononuclear-cells-pbm-cs-from-a-healthy-donor-v-1-0-1-1-standard-1-2-0 (atac_pbmc_1k_v1 dataset) and visualized using cellranger-atac-1.2.0 software.

### Generation of read density tracks

Homer v4.11.1 tools ^79^ were used to generate the tracks. First, bam files were converted to sam files using samtools v1.9 using the option “-G 1024” to use only unique reads at pseudo-bulk level. The sam files were processed with the makeTagDirectory() function to create tags for every chromosome and with makeUCSCfile() function to create BedGraph files. BedGraph files were then convefrted into the BigWig format using the bedGraphToBigWig v4 package in the collection of UCSC tools. BigWig files are suitable for uploading into the UCSC genome browser.

### Droplet quantifications by imaging

Three replicate emulsions (*n* = 3 experiments) were generated in 200 uL volumes in different days with an excess of DAPI-stained cells. Conditions fully replicated a standard AmpliDrop reaction except for the number of cells, which was much higher than usual to facilitate the finding of cell-encapsulating droplets under the microscope. A total of 142 images were taken from these emulsions using a 20x optical objective with 1.5x digital amplification (Keyance microscope, model: BZ-X710). The pictures were taken by moving the sample randomly until a cell-encapsulating droplet was detected by the operator. The picture was opened later with ImageJ v1.54d software and the diameter of the cell-encapsulating droplet was measured. A total of *n* = 200 cell-encapsulating droplets were detected and measured (blue line in Fig. 1b). An image from a Neubauer chamber (hemocytometer) taken with the same settings was used as reference for calibration of the ImageJ software. In addition, a total of *n* = 6 images were used for diameter quantification of all droplets, excluding those with a diameter of 10 microns or smaller. We note that, in the *n* = 142 images containing cell-encapsulating droplets, no cell was detected in a droplet with a diameter smaller than 10 microns, and only one cell was detected in a droplet with a diameter between 10 and 20 microns, while *n* = 199 cells were found in droplets of 20 microns or larger. These droplets represent 98.07% of the aqueous phase in the emulsion. In the six images used for diameter quantification of all droplets, we measured *n* = 802 droplets (green line in Fig. 1b). The measurements were also used to infer volumes (red line in Fig. 1b). We note that the actual measurements might not exactly represent the actual diameter of the droplets since the emulsions were placed in between two large coverslips under the microscope for proper focus and flattening of the solution. Nonetheless, the purpose of collecting these data was not to quantify the diameter of the droplets with precision, but to compare those with cell-encapsulating properties and those without cells, and to roughly estimate the number of droplets in an emulsion.

### Granular functional filtering (Gruffi) analysis

In R Studio (v2023.03.0+386) within a Seurat environment (v5.1.0), Gruffy analysis was performed using the Gruffi package v.0.7.4 ^50^ available at https://github.com/jn-goe/gruffi. We used the same parameters used for the generation of Seurat objects (nPCs = 30, dimensions = 1:20, reduction = “umap”). GO categories from hsapiens_gene_ensembl: GO:0006096 # Glycolysis; GO:0034976 # ER-stress; and GO:0042063 # Gliogenesis, negative filtering. Gruffi thresholds were selected using Shiny with a 90% quantile and proposed thresholds. Gruffi annotations (stressed and nostressed) were exported matching cellular barcodes and used as labels in projections plotted in Excel v.16.84 (Microsoft) obtained from Seurat-generated UMAP plots.

### Label transferring by reference mapping

In R Studio (v2023.03.0+386) within a Seurat environment (v5.1.0), reference mapping was performed according to https://satijalab.org/seurat/articles/multimodal_reference_mapping.html and using the reference atlas from https://atlas.fredhutch.org/data/nygc/multimodal/pbmc_multimodal.h5seurat ^34^. The AmpliDrop query was uploaded as a filtered-feature_bc_mtrix.h5 format from aggregated libraries. Parse Biosciences PBMC scRNA-seq data gene matrix dataset was obtained as described in https://support.parsebiosciences.com/hc/en-us/articles/360053078092-Seurat-Tutorial-65k-PBMCs from the resources.parsebiosciences.com/downloads site generated from a healthy donor (67,000 cells). Seurat objects were created with min.cells = 3 and min.features = 200, and SCTransform normalization. No further filtering was applied to fairly compare technologies without computational aid. The anchors were defined using the SCT normalization method and the spca reference.reduction argument with 50 dimensions. MapQuery used ADT as predicted_ADT and the reduction model wnn.umap. Image outputs were generated for predicted.celltype.l1 and predicted.celltype.l2 annotations. Outputs were generated with the DimPlot() function from the ggplot2 v.3.5.1 library.

### Manual labeling based on literature-supported gene markers

Manual cell annotations were performed using literature-searched markers and combining Seurat-defined clusters (generally at 0.5 resolution) accumulating most of the signal. When one cluster accumulated most of the expression signal of a marker, the cluster was labeled with the identity of the cells distinctively expressing the marker, based on the literature. When two or more clusters accumulated most of the expression signal of a marker, the clusters were combined to generate a larger cluster that was labeled with the identity of the cells distinctively expressing the marker, based on the literature. Cell line markers in mock mixtures were defined based on the literature or the AmpliDrop 3’ scRNA-seq analysis of the individual line: *ESR1, GREB1, BCL2* for MCF7 cells ^80^; *NRXN1, ASXL3, NLGN4X*, and *COL11A1* for A2780 cells and *EPHA2* for A2780cis cells ^28^; *EREG, CD44, PCDH7* for HCT-116, and *XIST, AKT3, CDH2* for 293T cells. PBMC markers were defined based on the literature: *PTPRC* for all PBMCs; *CD247* for all T and NK subtypes; *SLC8A1* for monocytes and cDCs; *AFF3* for B cells and the pDC and cDC populations; *MS4A1* for B cells; *BANK1* for B intermediate cells; *SSPN* for B memory cells; *COL19A1* for B naïve cells; *JCHAIN* for plasmablast; *SLC4A10* for MAIT; *RTKN2* for Treg cells; *VCAN* for CD14 classical monocytes; *TCF7L2* for CD16 non-classical monocytes; *NEGR1* for cDCm pDC, and HSPC populations; *CLNK* for cDC1; *NKAIN2* for HSPC; *UGCG* for pDC; *RBMS3* for pDC and Treg cells; *LDB2* and NCAM1 for NK_CD56bright cells; *GNLY* for NK cells; *CCL5* for CD8 TCM and proliferating NK cells; *IL7R* for CD8 TCM and others; *TSHZ2* for CD4 TCM and dividing, naïve, and dnT subtypes; *MKI67* for dividing populations of any type. Organoid markers included: *DCX* and *INA* for neuronal cells; *STMN2* for differentiated neurons; *MEIS2* for telencephalon cells; *RSPO2* for diencephalon cells; *LHX1* and *LHX5* for non-telencephalon cells; *VIM* for non-neuronal cells; *NEUROD2* and *BCL11B* for glutamatergic neurons (telencephalon); *EOMES* for glutamatergic IPCs; *HES6* for IPCs; *DLX6-AS1* and *GAD2* for GABAergic interneurons; *ERBB4* for migrating GABAergic interneurons; *NEFL* for early neurons; *RELN* for Cajal Retzius neurons; *ROBO1* for some GABAergic subtype; *SOX2* for progenitors; *MKI67* for dividing cells; *EGFR* for Pre-OPC; *TTYH1, SFRP1*, and *GLI3* for RG; *THSD4* for late RG; *IGFBP5* for ependymal and other ventricular cells; *TTC6* for ependymal cells; *TTR* and *HTR2C* for choroid plexus subtypes; *PCDH15* for oligodendrocytic lineage; *GFAP* for astrocytic lineage; and *CLU* and *BCAN* for astroglia.

### Gene expression analysis using the Single Cell Portal (Broad Institute)

To generate dot plots with gene expression values from 23-days, 1-month, 2-month, 3-month, and 6-month cortical organoids ^51^, we used the Cortical Organoids Atlas in the Single Cell Portal maintained by the Broad Institute available at https://singlecell.broadinstitute.org/single_cell/study/SCP1756/cortical-organoids-atlas. The genes included in the figures were all added to the search function in the Portal and the tab for Dot plot was used for visualization with default settings, without additional filtering. The Clustering option was changed to visualize the scRNA-seq data for each cell culturing time point using CellType for Annotation.

### Pseudo-bulk differential gene expression

In R Studio (v2023.03.0+386) within a Seurat environment (v5.1.0), differential gene expression was generated using the DESeq2 package (v.1.42.1) and a pseudo-bulk approach, which has been reported as more robust than single-cell-based approaches ^81^. The following labels were added to the Seurat object using the AddMetaData() function: AmpliDrop 3’ scRNA-seq libraries R1 and R2 and 10X 3’ scRNA-seq libraries R1 and R2, and processed using pseudo-bulk signal with the AggregateExpression() using the labels to compare AmpliDrop R1+R2 and 10X R1+R2 based on counts$RNA. Filters: rowSums(counts(dds)) >=10. Results were exported into csv file and imported for plotting using the EnhancedVolcano package (v.1.20.0), being color-coded in red and grey using pCutoff = 10e-32 and FCcutoff = |2|.

### DAVID GO analysis

Gene ontology (GO) analysis was performed using DAVID tools available at https://david.ncifcrf.gov/tools.jsp and supported by the DAVID Bioinformatics Team (LHRI/ADRD at Frederick National Laboratory) and funded by the National cancer Institute. Gene symbols were obtained from the csv output of the differential gene expression analysis comparing 10X and AmpliDrop libraries after sorting the data by padj and log2FoldChange to identify the genes with pCutoff < 10e-32 and FCcutoff > |2.0|, which were pasted separately (from the AmpliDrop and 10X lists) into the Enter Gene List window. The identifier selected was OFFICIAL_GENE_SYMBOL, the species selected was Homo Sapiens, and the list type selected was Gene List. The DAVID tool used was the Functional Annotation Tool and the Functional Annotation Chart using only symbols assigned to Homo sapiens. Results were then exported as a csv file and processed with Excel v.16.84 (Microsoft) by sorting the results by Category and FDR. The top twenty terms with the lowest FDR were selected within the GOTERM categories (GOTERM_MF_DIRECT, GOTERM_CC_DIRECT, and GOTERM_BP_DIRECT). The selected terms were plotted with Excel.

## Figures preparation

The figures were generated with PowerPoint v16.84.1 (Microsoft) and the art was created using Biorender.com licensed to I.G.B.

