## Supplemental Data for "Single-Molecule Barcoding Technology for Single-Cell Genomics"

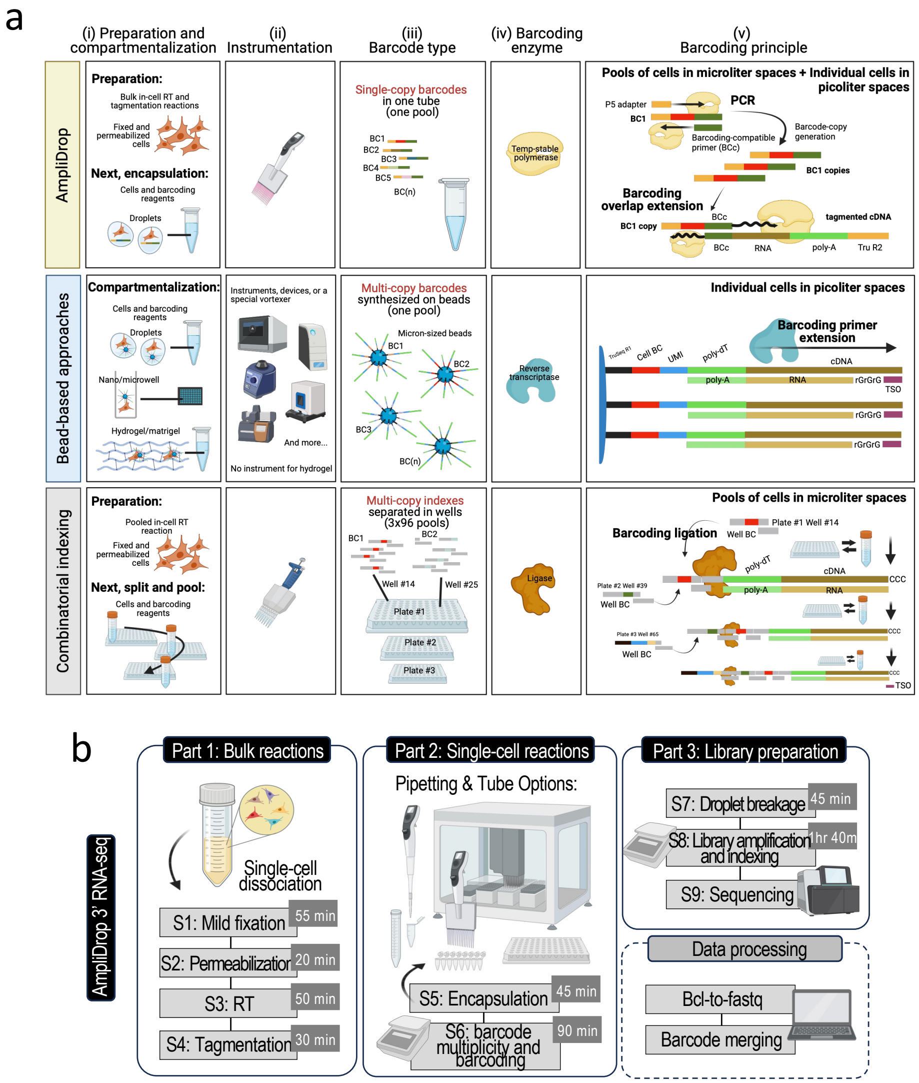


**Extended Data Fig. 1. AmpliDrop 3' scRNA-seq workflow and comparison with the rest of scRNA-seq barcoding principles. a**, Scheme depicting major differences between AmpliDrop and bead-based approaches—either droplet-based, microwell-based, or hydrogel-based methods for compartmentalization—and combinatorial indexing approaches. The major differences are organized into five sections: (i) differences related to cell preparation for barcoding and compartmentalization; (ii) differences related to instrumentation for compartmentalization; (iii) differences related to barcode type and moment of barcode multiplicity generation (during or prior library preparation); (iv) differences related to barcoding enzyme; and, (v) differences related to barcoding principle. The key elements introduced by AmpliDrop are: [1] in-cell bulk RT and tagmentation reactions, preparing cells for barcoding (these reactions must be preceded by fixation and permeabilization steps); [2] using a conventional electronic pipette to compartmentalize cell and the barcoding reactions; [3] using single-copy cellular barcodes, rather than multi-copy cellular barcodes, with the obligatory multiplicity generated during, rather than before, library preparation; [4] Using thermostable DNA polymerase to amplify single-copy barcodes and to introduce the amplified barcodes into cDNA though the shared BC'ing-compatible sequences (barcoding step). **b**, AmpliDrop barcoding technology applied to a 3' scRNA-seq workflow. The library preparation can be divided into three parts. Part 1 consists of bulk reactions: mild cell fixation and permeabilization, which enable in-cell preparation steps for barcoding, and RT and tagmentation reactions, which prepare cells for barcoding and requires synthesizing cDNA and incorporating BC'ing-compatible sequences. Part 2 consists of single-cell reactions: encapsulating cDNA-BC'ing-compatible cells with unique single-copy barcodes and barcoding/PCR reagents into water-in-oil droplets using an electronic or robotic system. Using a pipette provides high flexibility of tubes and volumes for barcoding, and a wide range of scales and manual/automation processes. Part 2 also includes the AmpliDrop reaction: first, barcode-multiplicity by PCR, then, barcoding reaction by overlap extension. The barcoding reactions relies on the same sequence present in the 3' end of the amplified barcodes and the 5' of the transposed cDNA molecules. And Part 3 consists in typical steps of library preparation, including droplet breakage, library amplification and indexing, and sequencing. Finally, data processing requires generating fastq files and barcode merging (see next sections for more details).

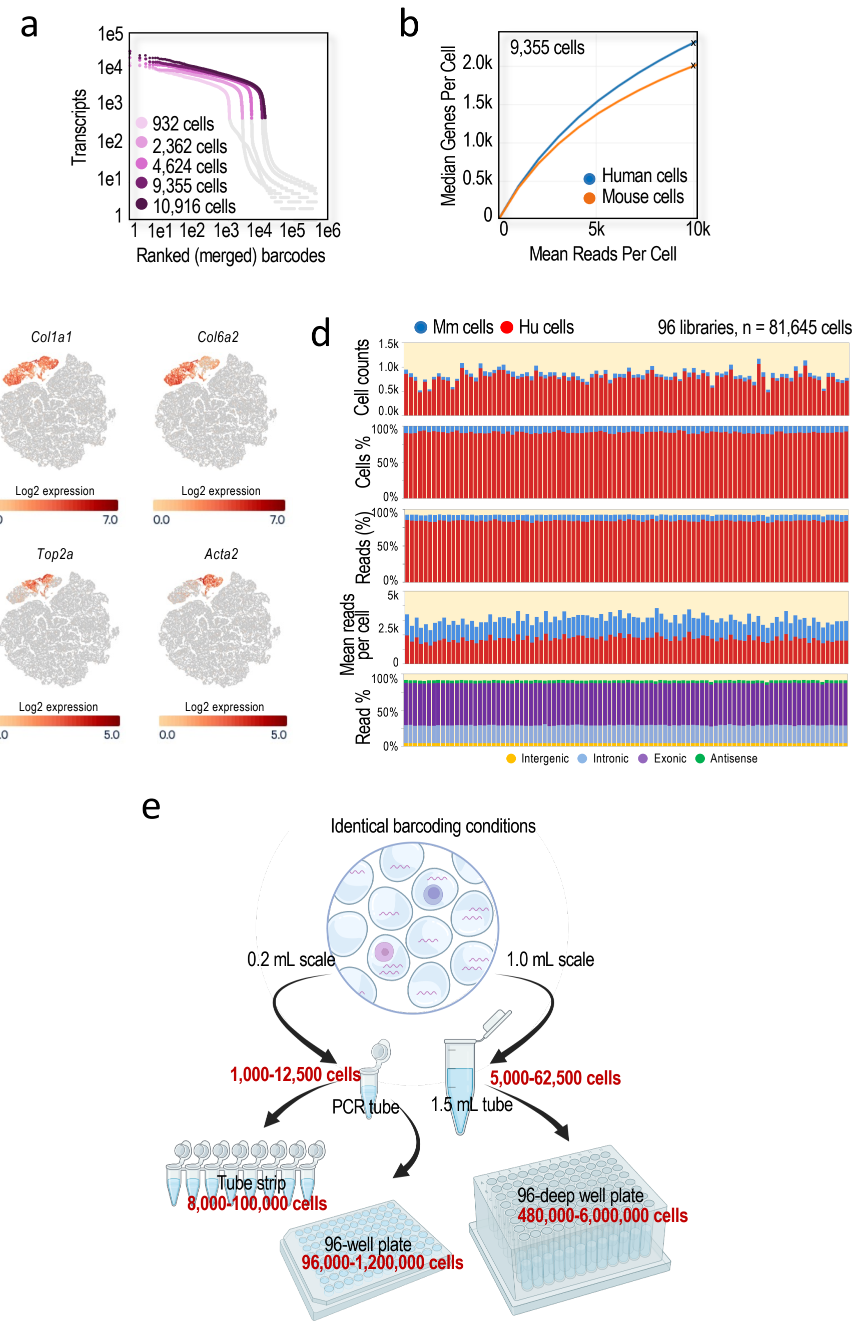


**Extended Data Fig. 2. Barnyard experiments and technology scalability**. **a**, Ranked (merged) barcode plot of AmpliDrop 3' scRNA-seq data generated from human and mouse mock mixtures listed in Fig. 1c. **b**, Plot of sequencing depth (mean reads per cell) relative to gene capture (median genes per cell) by species in the AmpliDrop 3' scRNA-seq experiment with an output of 9,355 human/mouse cells. **c**, Expression levels for the indicated mouse genes across the UMAP plot shown in Fig. 1g supporting sufficient resolution to distinguish some cell states. **d**, Data analysis for the experiment shown in Fig. 1g. (Top to bottom) Histogram of cell counts, cell proportions, read proportions, mean reads per cell, and genomic annotations by species and library. **e**, Results are shown in Fig. 1 validating the emulsifying conditions and droplet-to-cell and barcode-to-droplet ratios that can be similarly applied to a variety of tube formats and emulsion volumes, providing wide scalability: from 200 µL emulsions in a single PCR tube, a tube strip (8 or more), or 96-well plates, to 1 mL emulsions in single 1.5 mL tube or 96-deep-well plates. Thus, the same experimental conditions enable library preparation spanning from 1,000 cells in one single tube to, theoretically, 5-6 million cells on a plate.

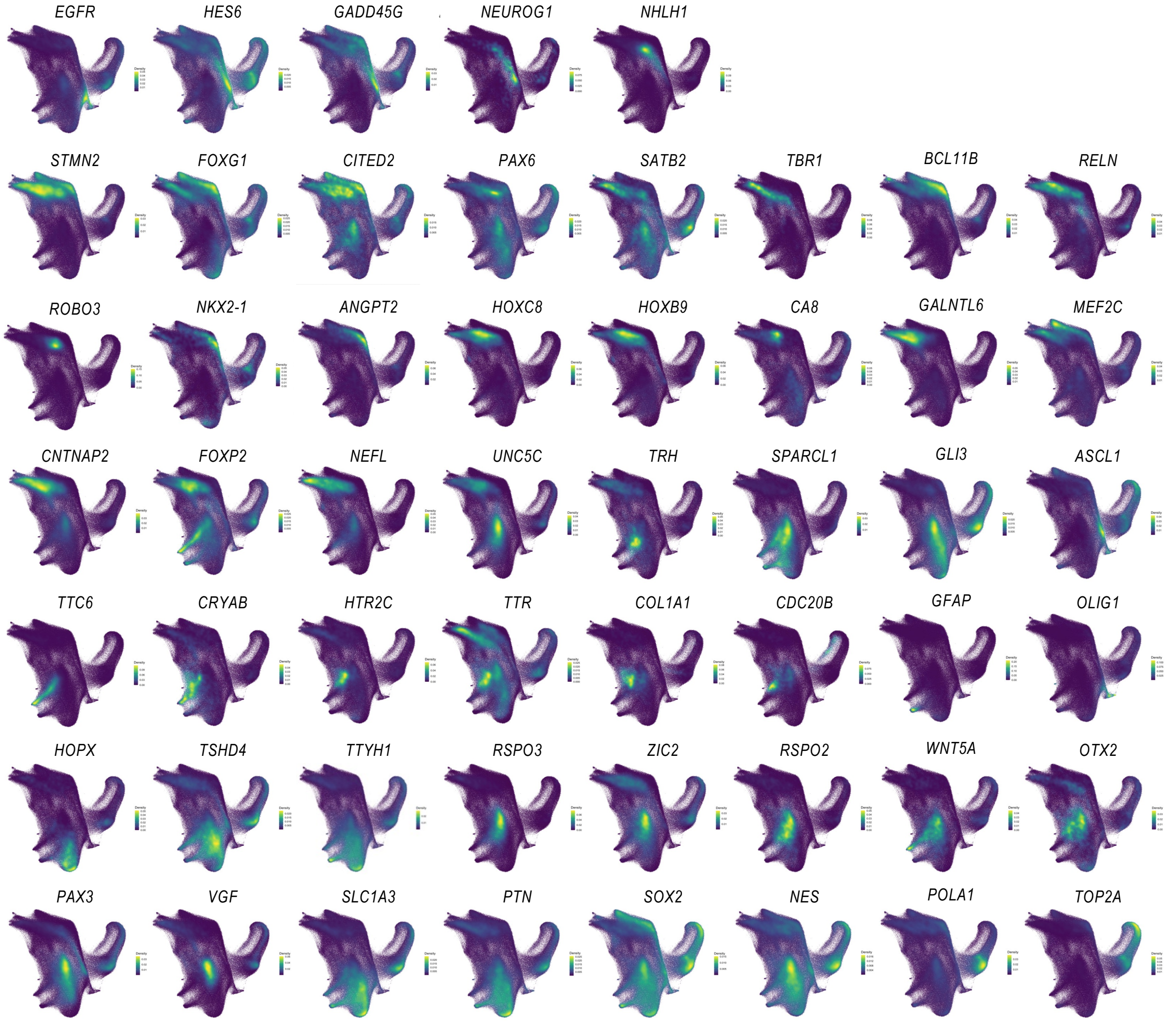


**Extended Data Fig. 3. Multi-library AmpliDrop 3' scRNA-seq integration**. AmpliDrop 3' scRNA-seq analysis of 72 libraries and almost six hundred thousand cells. Plots show count densities for the indicated genes. Cell annotations typically associated with the indicated genes have also been included.

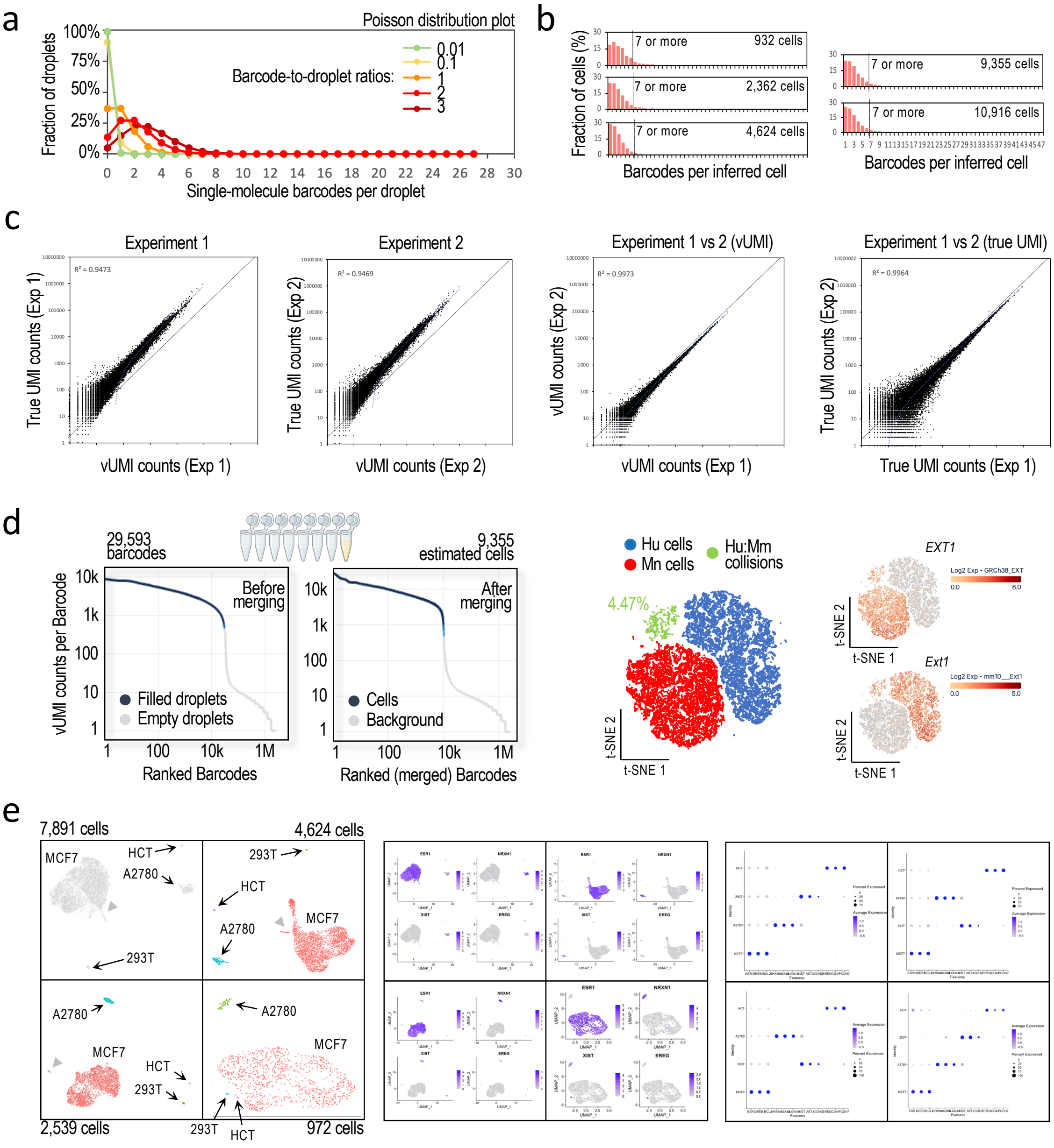


**Extended Data Fig. 4. Barcode merging to infer cells**. **a**, Predicted percentage of droplets (y-axis) containing the indicated number of single-copy barcodes (x-axis) according to a Poisson distribution and barcode-to-droplet ratio (color-coded lines). For instance, a barcode-to-droplet ratio of 0.01 is expected to generate only one barcode-containing droplets, but this condition will result in more than 99% of barcode-empty droplets. On the other hand, a barcode-to-droplet ratio of 3 will result in less than 10% empty droplets, with the caveat that most barcode-containing droplets are expected to contain more than one barcode (multi-barcoded droplets). If the goal is minimizing cell dropouts, a barcode-to-droplet ratio of 3 or more is necessary, which will require a computational tool to identify multi-barcoded instances. **b**, Inferred distribution of non-barcoded, single-barcoded, and multi-barcode cells in the experiment shown in **Fig. 1c**, as labeled. In these experiments, most cells are inferred as multi-barcoded (70%), with a few based on more than six barcodes per droplet, which eliminate during data analysis, in part because these cases likely represent collisions. **c**, Pairwise comparisons of vUMI and true UMI counts as indicated. **d**, (*Left panels*) Ranked barcode plots pre and post merging in the barnyard experiment with an output of 9,355 human/mouse cells (compare to t-SNE plots shown in **Fig. 1e** based on merged data). (*Right panels*) t-SNE plots with cells color-coded by read mapping preference: mostly mapping to human in blue and mouse genomes in red. Cells considered as human-mouse cell collisions are shown in green. Annotations based on the expression of human and mouse genes (e.g., human *EXT1*- and mouse *Ext1*-expressing cells; expression levels are shown in log2 scale), and human-mouse cell collisions are inferred by their separate clustering. **e**, UMAP plots of AmpliDrop 3' scRNA-seq data from the MCF-7, A2780, 293T, HCT-116 mix shown in **Fig. 2a**. (*Left*) Clusters were labeled and color-coded based on markers identified in single-cell line experiments. Cell outputs are indicated (with the plot based on $n$ = 7,891 cells repeated here from **Fig. 2a**; sequencing depth, $n$ = 11,981 (972 cells), 14,285 (2,539 cells), 12,986 (4,624 cells), and 14,090 (7,891 cells) mean reads per cell). Note: a sub-cluster of MCF7 cells is highlighted with grey arrowheads in the three plots with the largest cell numbers. (*Center*) Expression labels of line-specific markers across the UMAP plots. (*Right*) Expression labels of line-specific markers in dot-plot format by inferred cell line.

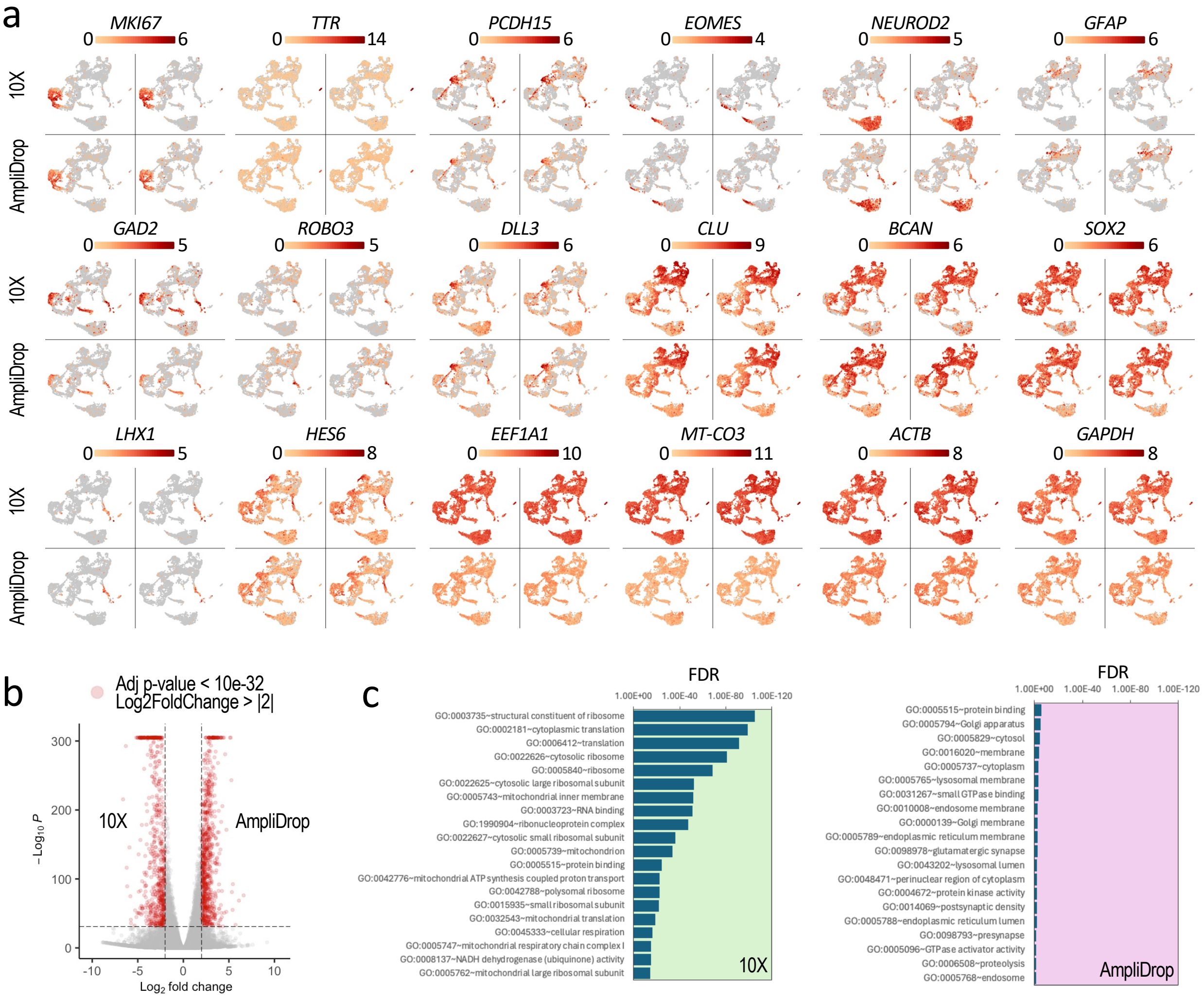


**Extended Data Fig. 5. Benchmarking AmpliDrop 3' scRNA-seq with the 10X Genomics 3' scRNA-seq technology. a**, Log2 expression levels of brain cell markers depicted on the UMAP plot integrating the two AmpliDrop and two 10X replicate libraries from the same sample shown in Fig. 3a but separated by replicate and single-cell technology, as indicated. **b**, Enhanced Volcano plot of pseudo-bulk differential gene expression between AmpliDrop and 10X Genomics libraries ($n$ = 4 libraries, $n$ = 28,331 variables). The top differentially expressed genes with significantly adjusted $p$-values < 10e-32 and Log2FoldChange > |2| enriched in the 10X experiment (left side of the plot) and the AmpliDrop experiment (right side of the plot) were selected and labeled in red ($n$ = 685 and $n$ = 870, respectively). **c**, Top twenty DAVID-enriched GO terms separated by technology in the most differentially expressed genes labeled in red in b. The x-axis represents significantly adjusted $p$-values (FDR) values. Plots show that 10X data is highly enriched in ribosomal, RNA metabolism, and mitochondrial terms, whereas AmpliDrop data shows not robust enrichment in any particular GO term.

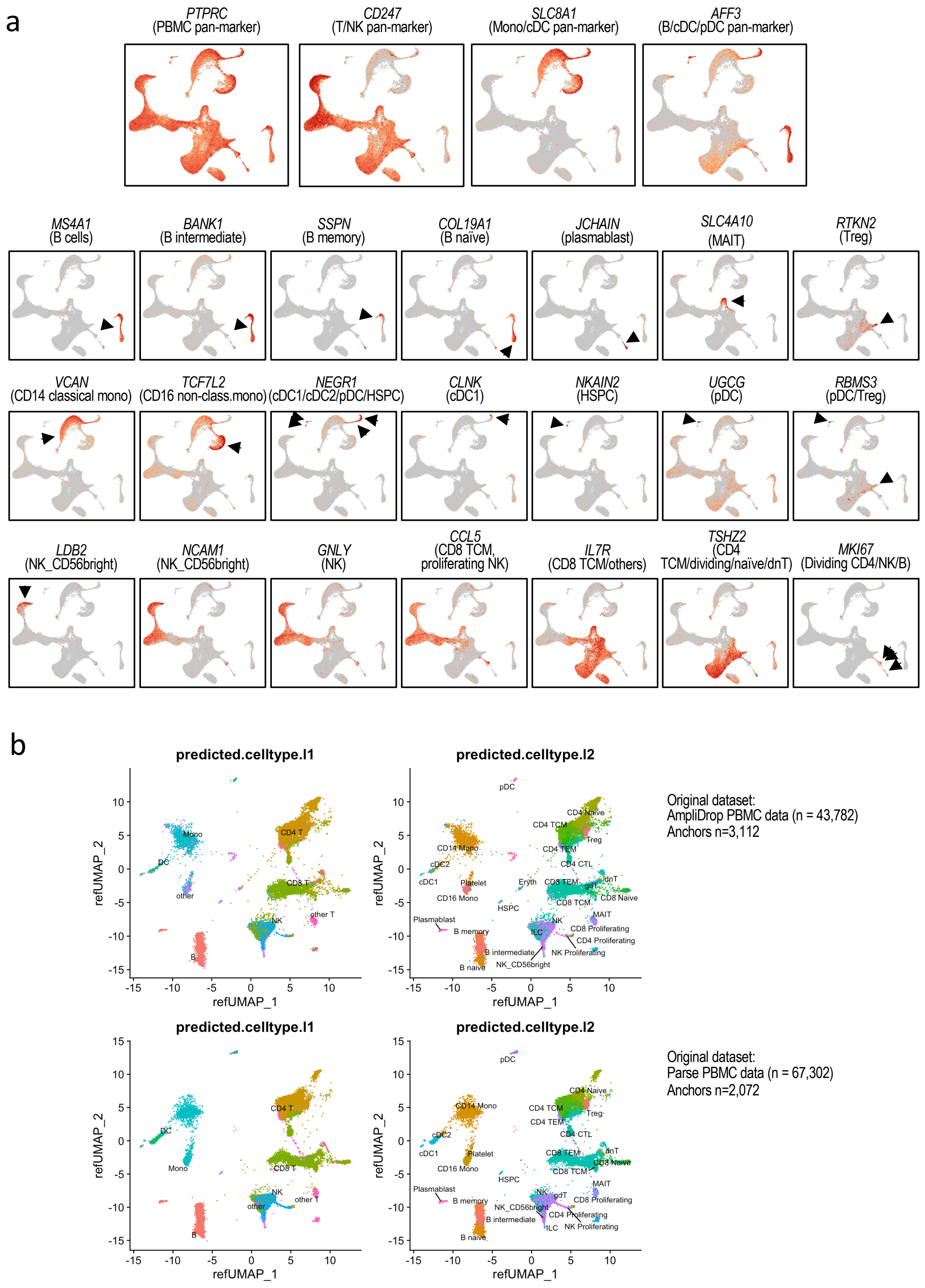


**Extended Data Fig. 6. Benchmarking AmpliDrop 3’ scRNA-seq with Parse scRNA-seq data. a**, Expression levels of well-known PBMC markers on the UMAP plots shown in Fig. 3c. The associated cell identities are also indicated. For small cell populations, arrowheads help to locate the annotated cells. **b**, Comparative analysis of label transfers over AmpliDrop and publicly available Parse PBMC scRNA-seq data from the same multimodal reference cell atlas [32].

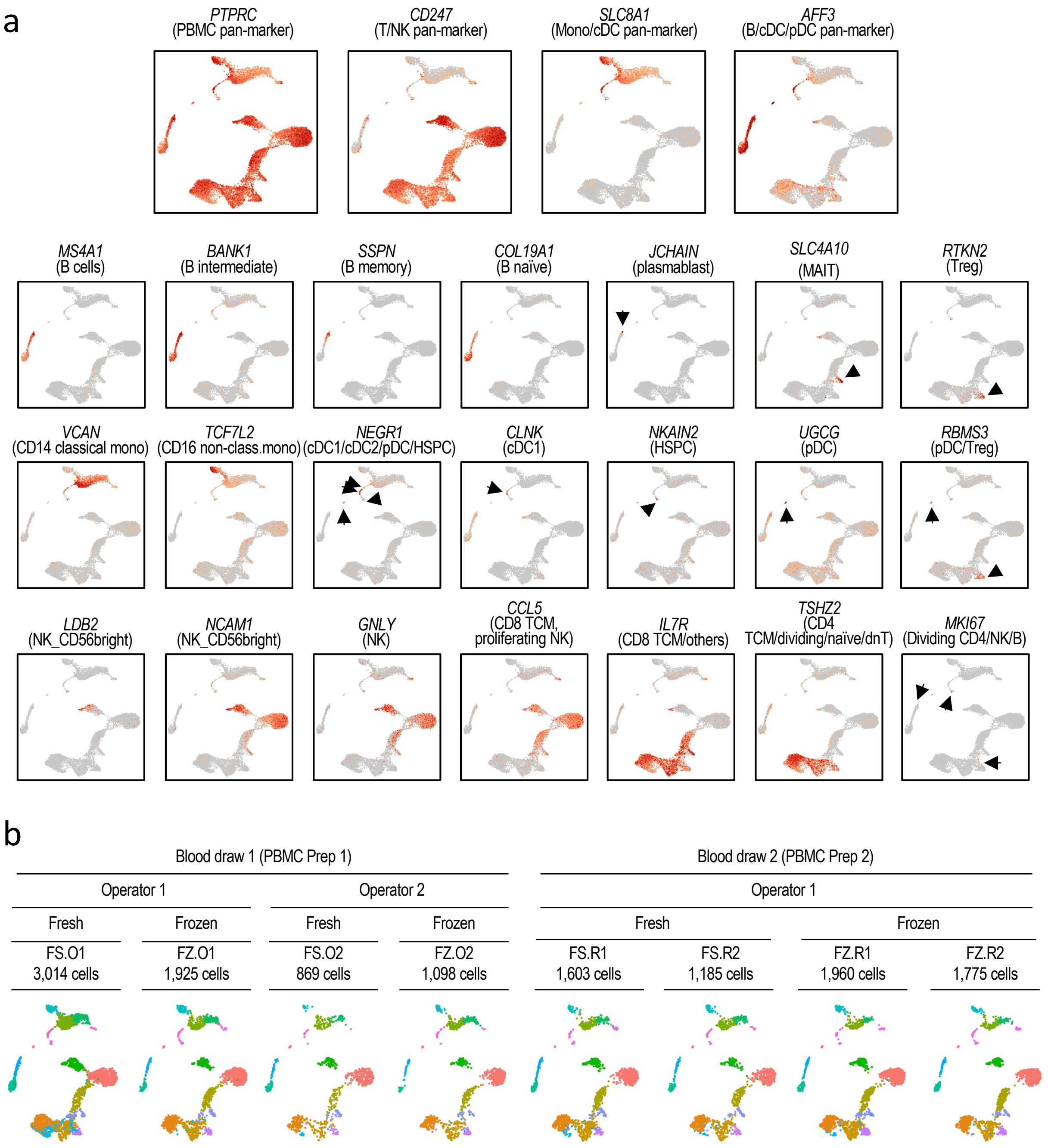


**Extended Data Fig. 7. Assessing the robustness of the AmpliDrop 3' scRNA-seq assay with PBMCs. a**, Expression levels of well-known PBMC markers on the UMAP plot shown in Fig. 3d. The associated cell identities are also indicated. **b**, UMAP plots shown in Fig. 3d separated by condition here.

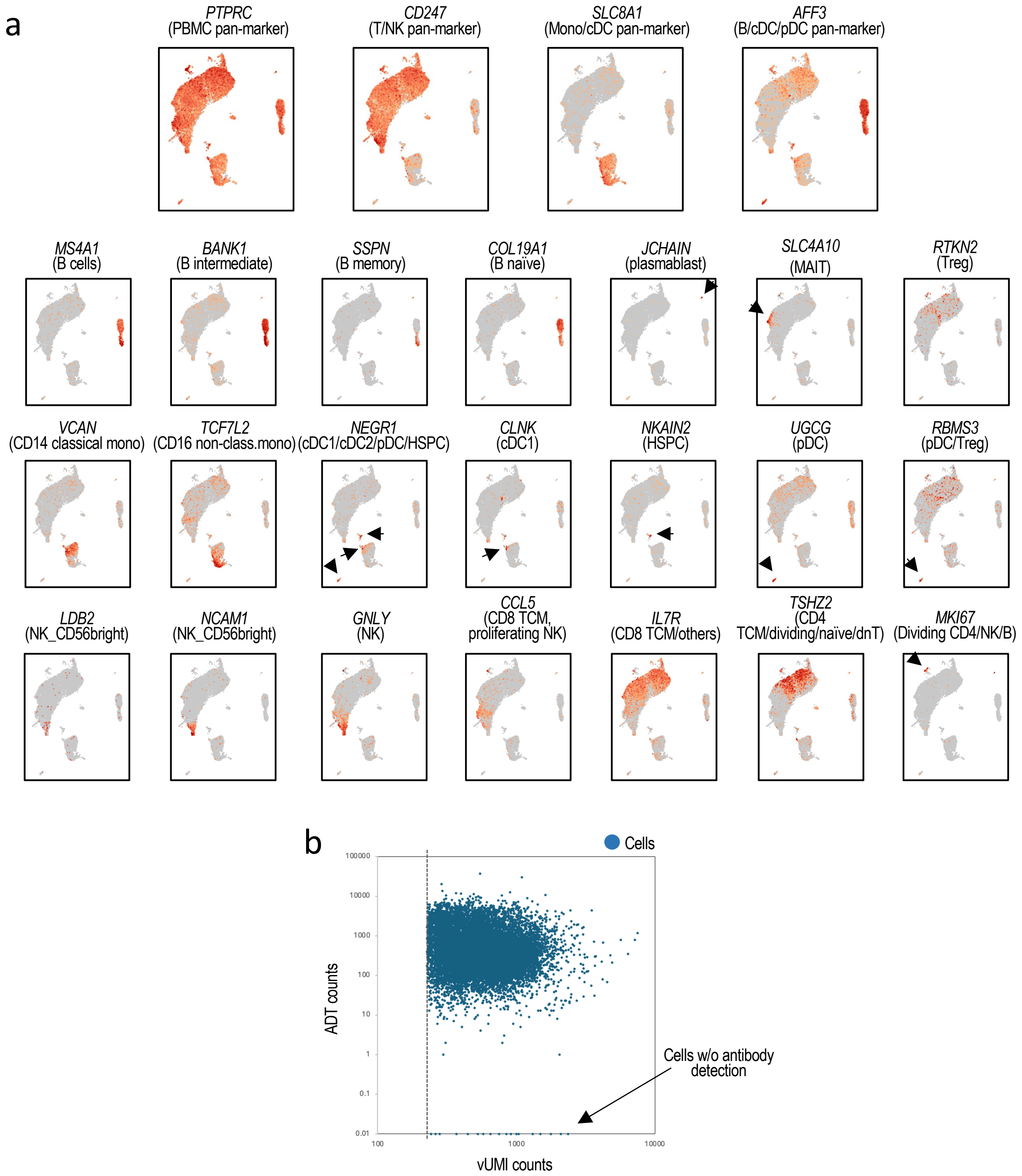


**Extended Data Fig. 8. AmpliDrop CITE-seq analysis with PBMCs. a**, Expression levels of well-known PBMC markers on the UMAP plot shown in Fig. 3e. The associated cell identities are also indicated. **b**, Scatter plot shows cells organized by total vUMI counts relative to total ADT counts. Virtually all cells have some ADT signal, except a few cases pointed by the arrow. We note that the used antibody cocktail, not only contain antibodies against all major immune cells, but also against CD45, which is expressed by all PBMCs.

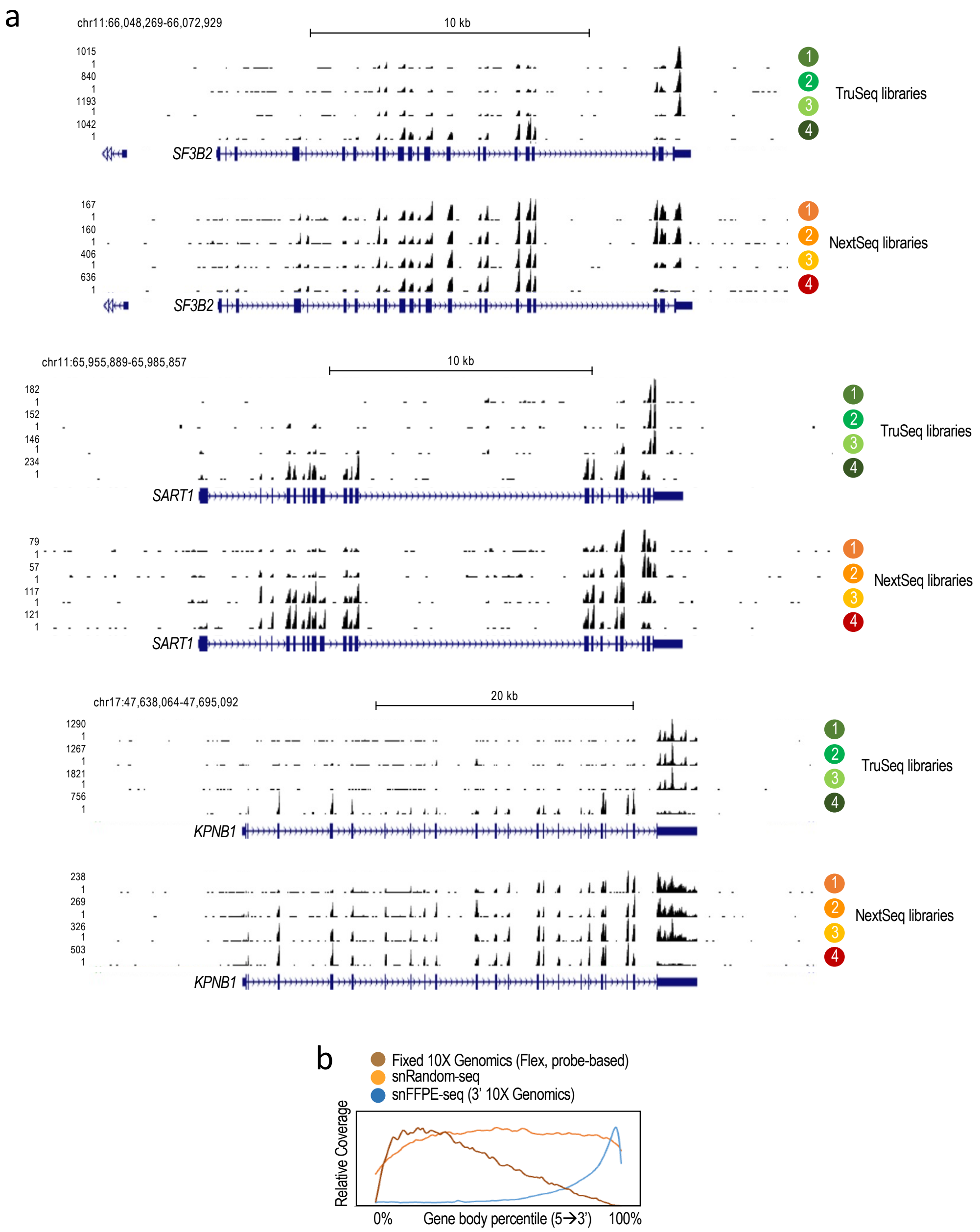


**Extended Data Fig. 9. AmpliDrop full-length scRNA-seq analysis. a**, AmpliDrop scRNA-seq read densities across the indicated loci matching the numbered AmpliDrop strategies depicted in the scheme shown in Fig. 4a. TruSeq libraries shown on top and NextSeq libraries shown at the bottom for each locus, as indicated. **b**, Normalized meta-profiles of signal across gene bodies for the listed technologies based on previously published data, used here as reference.

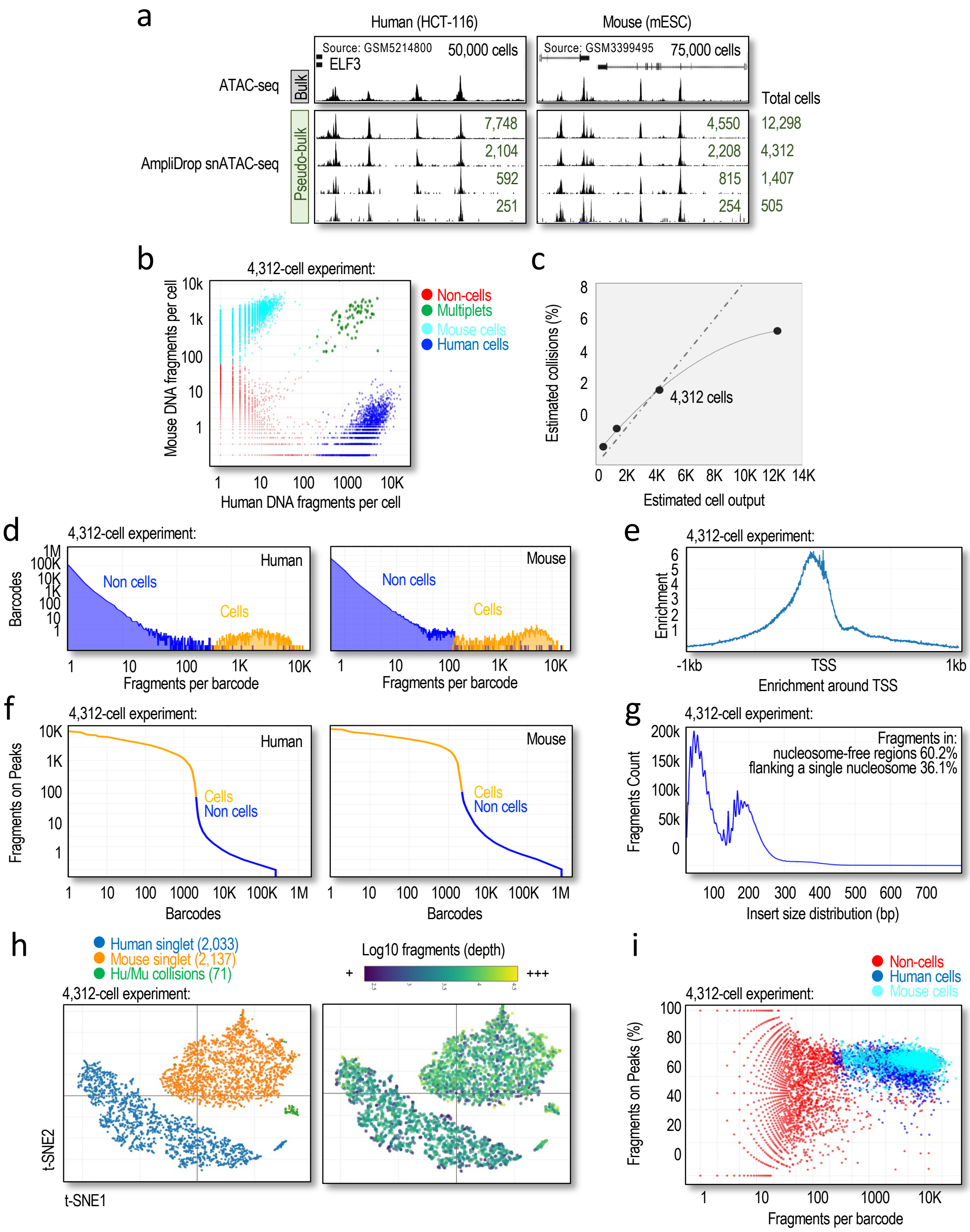


**Extended Data Fig. 10. AmpliDrop snATAC-seq analysis. a**, Bulk (*top*) and pseudo-bulk (*bottom*) read densities across the indicated loci showing data similarity between publicly available bulk ATAC-seq data (GSM5214800 and GSM3399495) and AmpliDrop snATAC-seq data (generated here) based on human HCT-116 cells (*left tracks*) and mESCs (*right tracks*). Input cells for the bulk experiments: $n$ = 50,000 and $n$ = 75,000, respectively. Output cells for the single-cell experiments (*top to bottom*): 12,298, 4,312, 1,407, and 505 cells (sequencing depths, $n$ = 11,504-18401 mean fragments per cell). **b**, Log-scale scatter plot of human (x-axis) and mouse (y-axis) DNA fragments per cell from the AmpliDrop snATAC-seq experiment with a total cell output of 4,312 cells in a. **c**, Collision rate estimates for the experiments shown in a. **d**, Barcode abundance by amount of DNA fragments in the library with an output of $n$ = 4,312 cells. **e**, Total read distribution around TSS in the library with an output of $n$ = 4,312 cells showing enrichment upstream of TSS. **f**, Ranked (merged) barcodes plot based on estimated fragments on peaks in the library with an output of $n$ = 4,312 cells. **g**, Fragment

(see next page)

size distribution shows a nucleosomal-like pattern of tagmentation in the library with an output of *n* = 4,312 cells. **h**, t-SNE plots from the library with an output of *n* = 4,312 cells showing human and mouse cell separation. **i**, Fragments per barcode relative to fragments on peaks color-coded by cell calling: human cells, mouse cells, and non-cells showing signal on peaks dominates when cells are called in the library with an output of *n* = 4,312 cells.

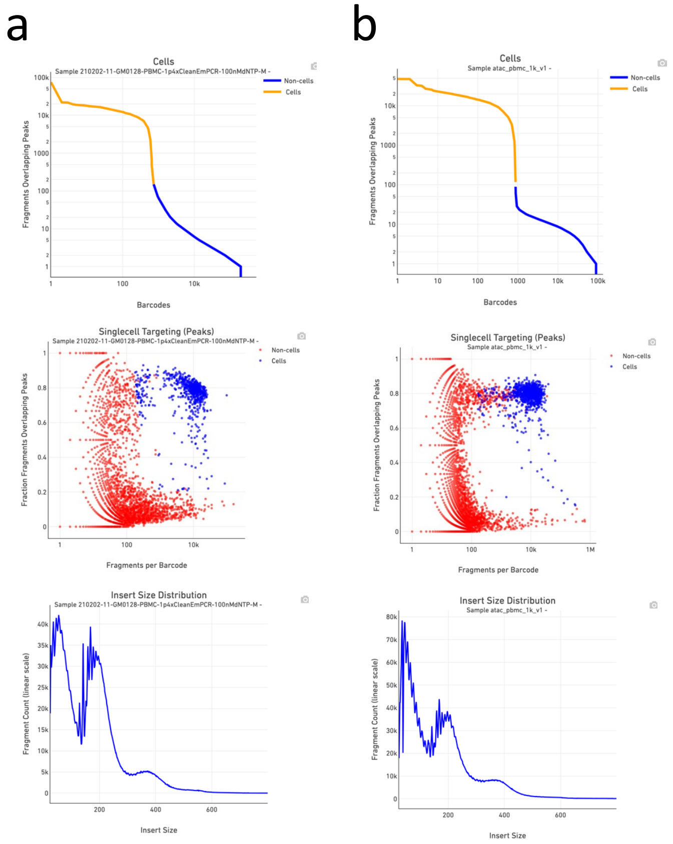


**Extended Data Fig. 11. Benchmarking AmpliDrop snATAC-seq with 10X Genomics snATAC-seq data a,b**, AmpliDrop (in a) and 10X Genomics (in b) snATAC-seq performance plots for the data shown in Fig. 4b,c. (*Top*) Ranked (merged) barcodes plot based on estimated fragments on peaks. (*Center*) Fragments per (merged) barcode relative to fragments on peaks. (*Bottom*) Fragment size distribution shows a similar nucleosomal-like pattern of tagmentation in both libraries.

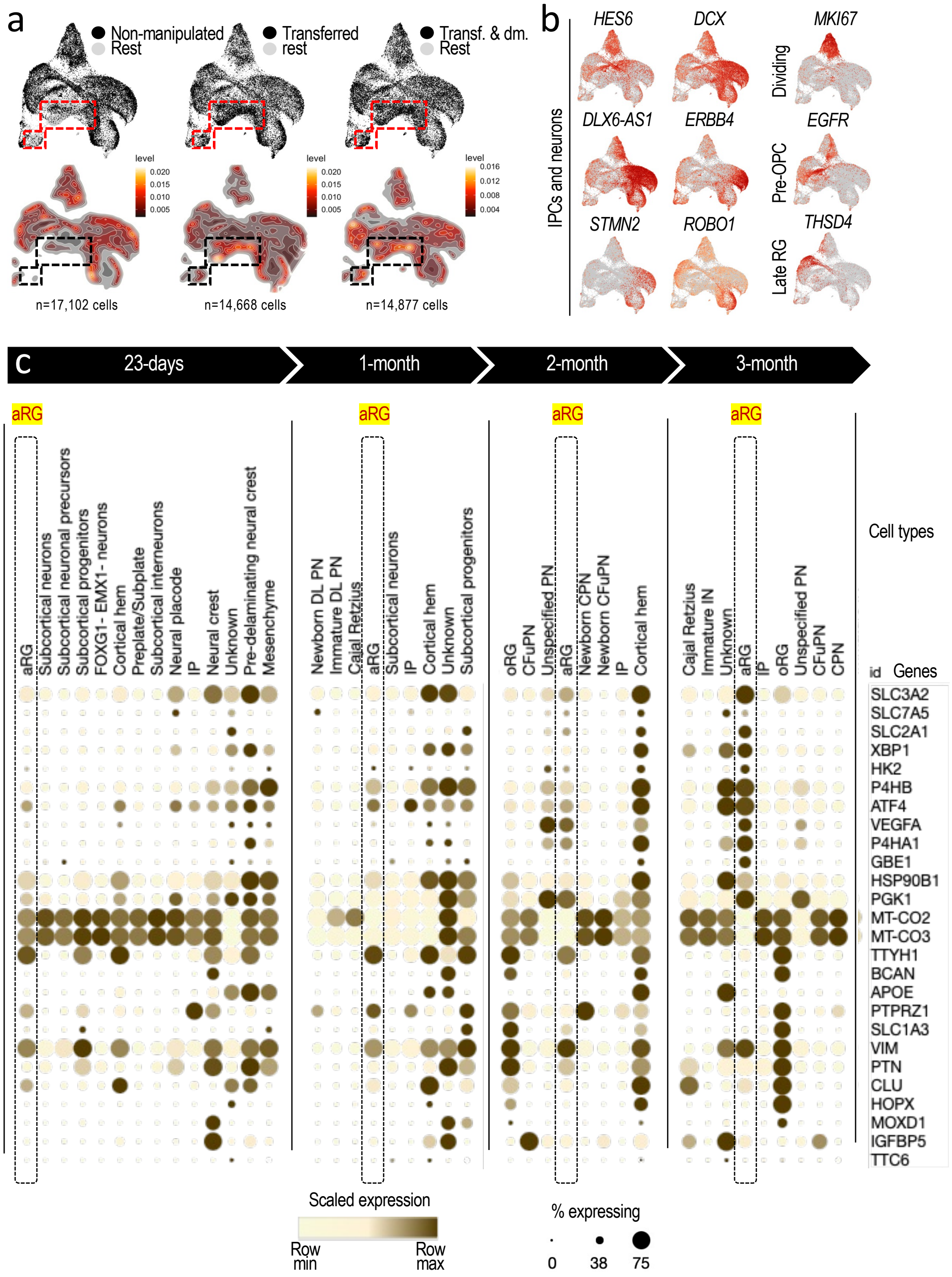


**Extended Data Fig. 12. Profiling the effects of physical manipulation on human cortical organoids.** **a**, (*Top*) UMAP versions of the plot shown in Fig. 5b highlighting cells (in back) by condition, as indicated. (*Bottom*) Cell density versions of the plots shown on top highlighting the areas with the highest cell density in every condition. **b**, Expression levels of well-known brain cell markers on the UMAP plot shown in Fig. 5b with cell identity annotations. **c**, Relative expression of the indicated genes generated with the Single Cell Portal from the Broad Institute and organoid data (23-days, 1-month, 2-months, and 3-months) and cell annotations from Ref. [45]. Gene selection as in **Fig. 5e**. Dotted rectangles highlight signal for aRG cells in every timepoint. A 6-month timepoint is shown in Fig. 5e. Notice the glycolytic nature of the aRG-like population gradually evolves with the age of the organoid. We suspect that the "unknown" population in the 3-month timepoint might represent a tRG-like version based on the expression of the ventricular *IGFBP5* gene.

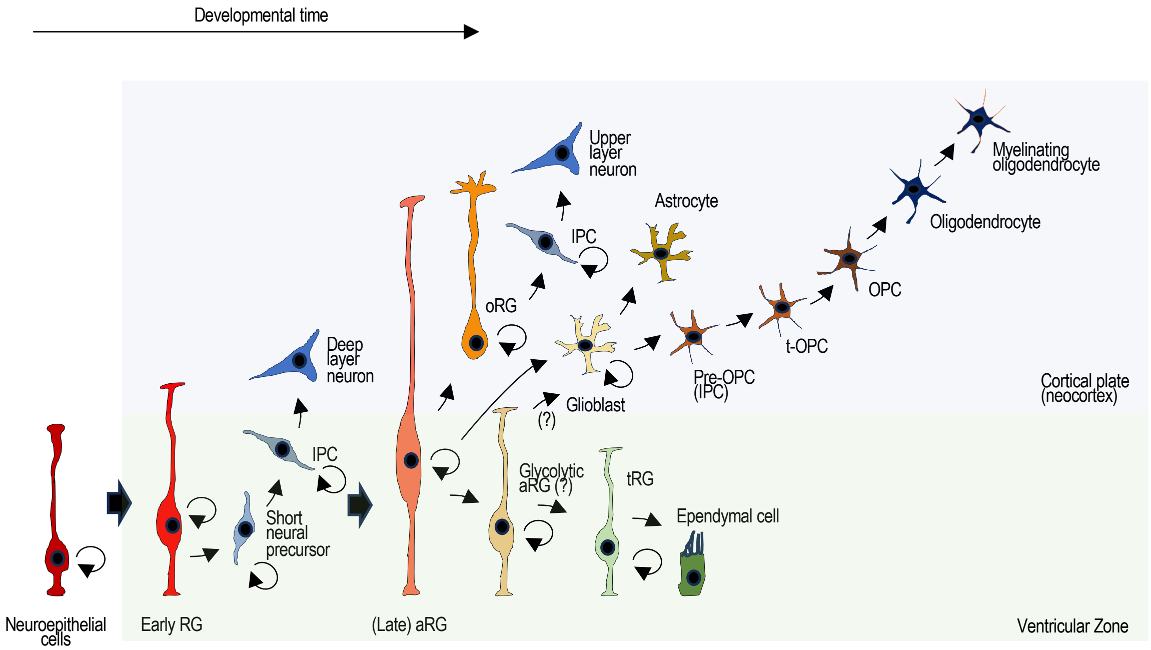


**Extended Data Fig. 13. Major cell types during early human cortical development.** Illustration depicting the major cell types in human early cortical development (adapted from Ref. [47]). Early during embryonic development, neuroepithelial cells differentiate into early RG, which are neural stem cells that further differentiate into early neuronal lineages (short neural precursor → intermediate progenitor cell or IPC → deep layer neuron) or proliferate (stemness). Later during embryonic development, early RG differentiate into aRG, which can further differentiate into outer RG (oRG) and truncated RG (tRG) depending whether the cells migrate into the cortical plate or remain within the ventricular zone, respectively. Outer RG can give rise to IPCs and a diversity of neuronal types. Truncated RG (cluster 6) can give rise to ependymal cells. Based on our observations, we propose that the glycolytic subtype of aRG (cluster 5) might be a precursor of tRG or, perhaps, a precursor of glioblasts with distinct differentiation properties. Pre-OPC (oligodendrocyte precursor cell), t-OPC (transitioning OPC, based on Ref. 69.

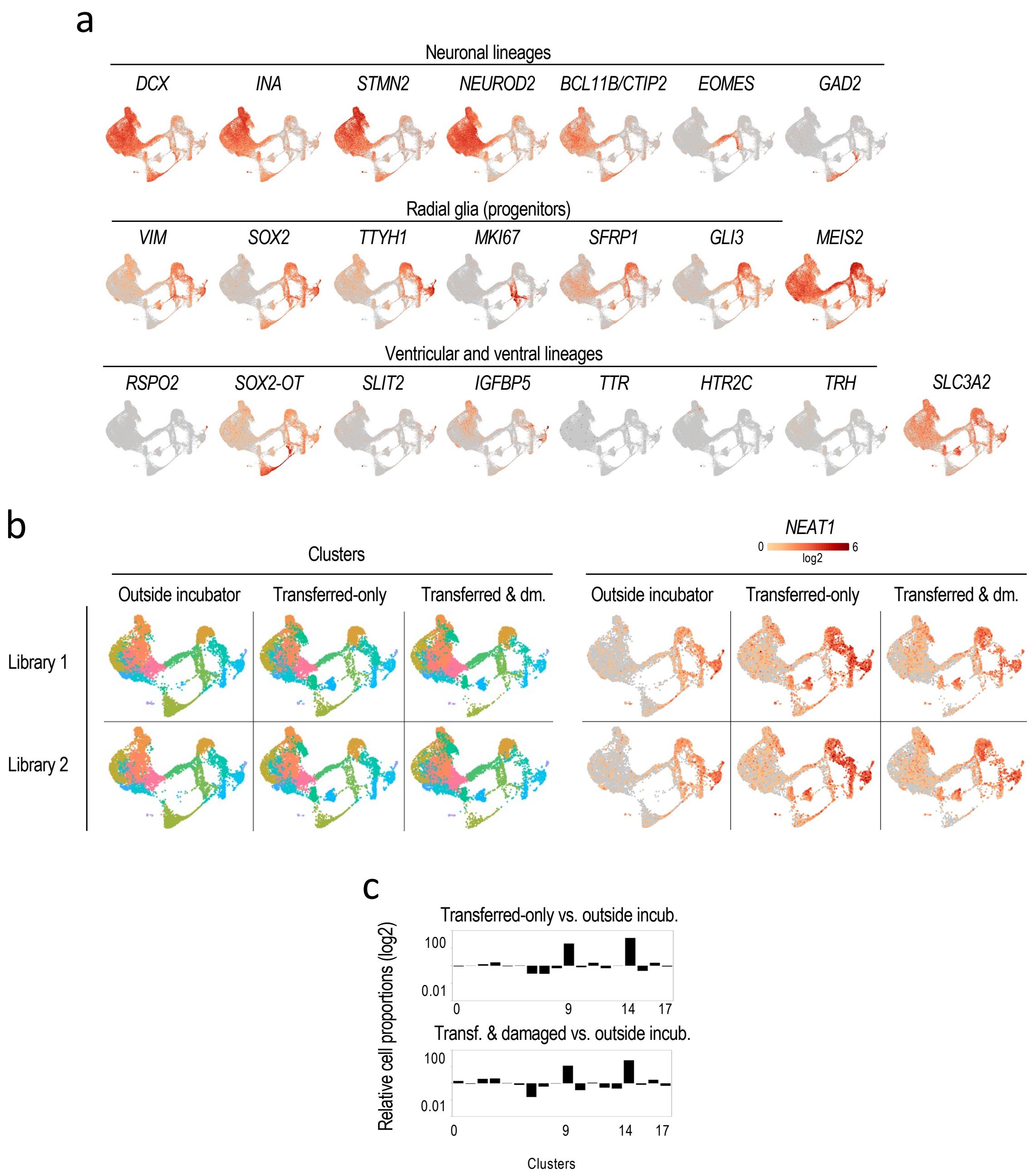


**Extended Data Fig. 14. Comparative analysis of cell composition among three sets of 6-month-old forebrain organoids exposed to mild and severe physical perturbations and unperturbed, with all sets maintained outside the incubator for approximately the same period (approximately 90 min). a**, Expression levels (log2 scale) of well-known brain cell markers on the UMAP plot shown in **Fig. 5g** with cell identity annotations on top of the gene symbols, as indicated. **b**, (*Left, clusters*) Individual UMAP plots separated by condition and library from the integrated analysis shown in **Fig. 5g**. (*Right, NEAT1*) Same plots shown on the left depicting *NEAT1* expression levels (log2 scale). **c**, Relative proportions (log2-scale ratios) by cluster (clusters shown in **Fig. 5g**) between non-perturbed and mildly physically perturbed (*top*) and severely physically perturbed (*bottom*) organoids.

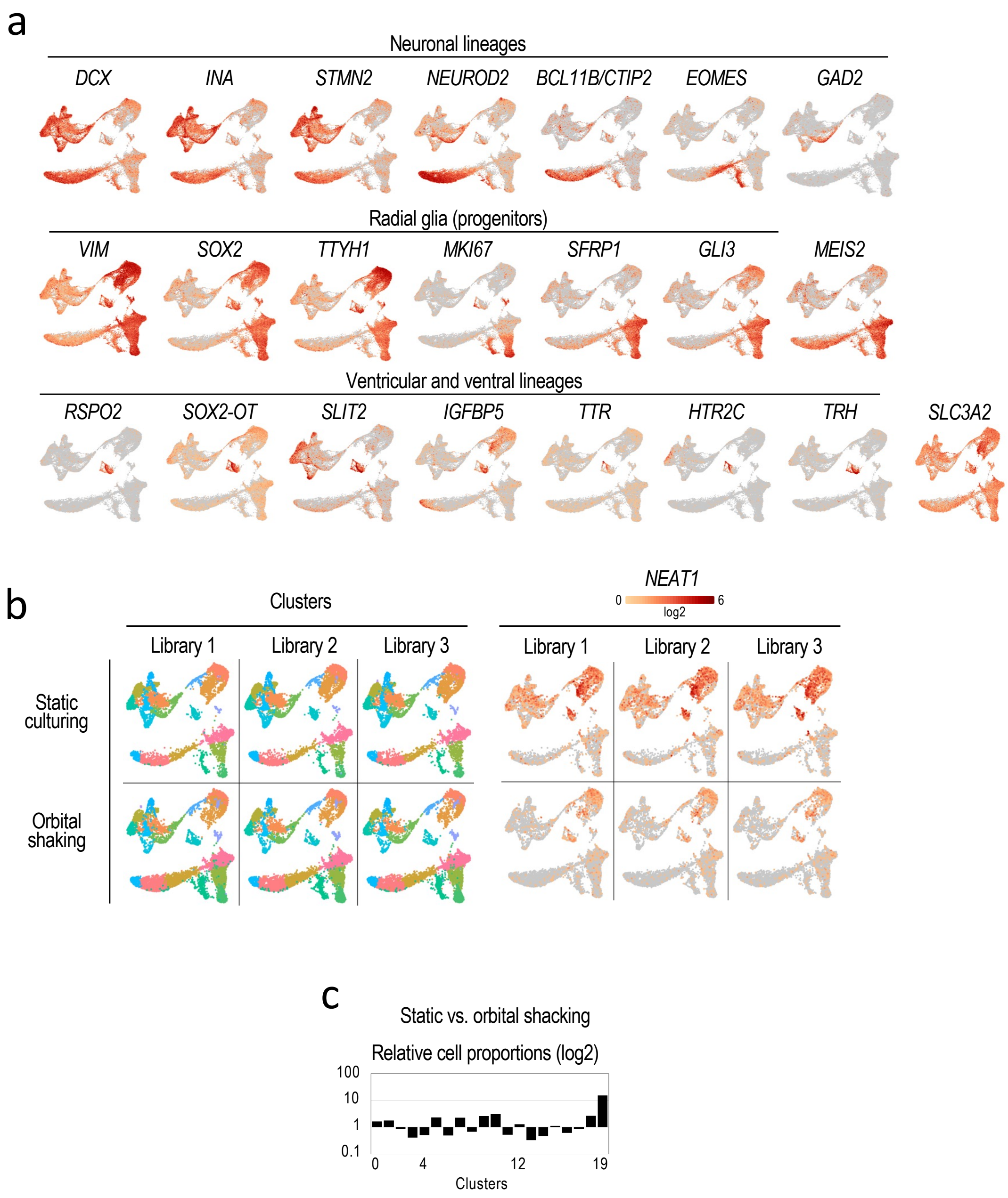


**Extended Data Fig. 15. Comparative analysis of cell composition between two sets of 2-month-old forebrain organoids cultured with or without orbital shaking (static) for 3 days prior to AmpliDrop library preparation. a**, Expression levels (log2 scale) of well-known brain cell markers on the UMAP plot shown in **Fig. 5h** with cell identity annotations on top of the gene symbols, as indicated. **b**, (*Left, clusters*) Individual UMAP plots separated by condition and library from the integrated analysis shown in **Fig. 5h**. (*Right, NEAT1*) Same plots shown on the left depicting *NEAT1* expression levels (log2 scale). **c**, Relative proportions (log2-scale ratios) by cluster (clusters shown in **Fig. 5h**) between 3-day orbital shacking and static culturing.
